# A plant-centric investigation of Class B Flavin-dependent Monooxygenase evolution and structural diversity

**DOI:** 10.1101/2025.09.16.676513

**Authors:** Joachim Møller Christensen, Elizabeth HJ Neilson

**Author notes:** Author contributions: JMC wrote this manuscript with input from EHJN.

## Abstract

Class B Flavin-dependent monooxygenases (FMOs) are ancient and ubiquitous enzymes, present throughout diverse kingdoms of life. These enzymes utilize flavin-based coenzymes to incorporate an oxygen atom into their substrate, thereby altering its chemical properties. In plants, FMOs perform crucial catalytic functions essential for plant health, particularly within hormone biosynthesis, defense compound production, and immunogenic responses. Despite the evolutionary significance of class B FMOs across life, their evolution within the plant lineage remains underexplored and underrepresented. Here we present a comprehensive, plant-centric phylogenetic investigation of class B FMOs to uncover lineage-specific evolutionary patterns and structural diversification. Using known Class B FMOs as baits, a large selection of flavin-related proteins was assembled from species representing key lineages across Viridiplantae. Structural domain architecture and motif analysis was used to accurately define different Class B FMOs, resulting in eight distinct class B FMO families. Three families include the canonical YUCCA, N-OX and S-OX FMOs, which remain the most abundant and prevalent across the plant kingdom. Three families are novel, encompassing a small selection of bryophyte FMOs. The analysis also expanded the BVMO family to include monilophyte and angiosperm members, and a potential YUCCA-related family is reclassified as the evolutionary distinct “Seedless FMO” family. Considerable structural diversification within the NADPH-binding domain is observed across the eight families, and by assessing structurally conserved folds rather than amino acid sequence, a refined set of conserved FMO-specific motifs is defined. Overall, this phylogenetic and structural analysis provides new insights into FMO evolution and provides an important foundational framework to aid functional characterization of class B FMOs in plants.

## Introduction

Flavin-dependent monooxygenases (FMOs) are ancient enzymes widespread through different kingdoms of life. FMOs utilize flavin-based coenzymes to incorporate an oxygen atom into their substrate altering its chemical properties such as bioactivity and solubility. FMOs are therefore essential for metabolic processes such as xenobiotic detoxification, while also possessing relevant properties for applications as industrial biocatalysts. FMOs have been classified into eight distinct classes (A–H) based on structural and functional characteristics (Huijbers et al., 2014; Mascotti et al., 2016)(Figure 1). Classes A, B, E, F and G are proposed to share a common evolutionary ancestor, with evolutionary divergence resulting from recruitment of different domains (Mascotti et al 2016). Some classes (A, B and G) function as single-component enzymes, while Classes C–F require a reduction partner for activity (Mascotti et al., 2016).

**Figure 1.**
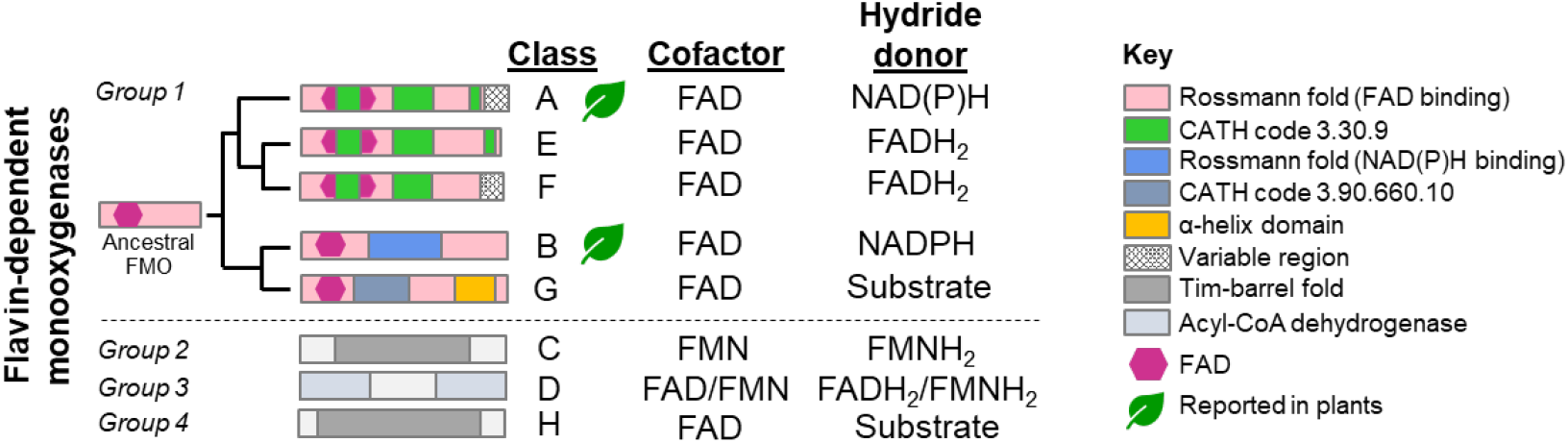
Schematic overview of flavin-dependent monooxygenase (FMO) classes with different flavin co-factors and hydride donors. FMOs are separated into eight classes, where classes A, B, E, F and G share a common evolutionary ancestor and FAD binding structural similarities. Plants possess FMOs belonging to classes A and B. Figure adapted from Mascotti et al., 2016.

Among the plant kingdom, class B FMOs are the most prevalent, abundant and diverse; present in all plant species, and in high copy numbers compared to other organisms (Thodberg and Neilson 2020). In fact, outside class B, only a single FMO has been characterized: a class A FMO involved in nitrogenous volatile compound biosynthesis in tomato (*Solanum lycopersicum*; Liscombe et al., 2022). Class B FMOs possess a distinct and characteristic domain architecture, characterized by two dinucleotide-binding domains. This structure arises from a duplication and subsequent functionalization of the FAD-binding domain (CATH 3.50.50.60) into an NADPH-binding domain, which results in the FAD-binding domain being split into two non-contiguous segments (**Figure 1**; Mascotti et al., 2016). Correspondingly, class B FMOs possess two conserved rossmann fold motifs important for the binding of the two di-nucleotides, FAD (GxGxxG) and NADPH (GxGxxG/A). Class Bs also possess a so-called “FMO identifying” motif (FxGxxxHxxxY/W), where the tryptophan is unique to a clade of class B FMOs known as Baeyer-Villiger monooxygenases (BVMOs), that catalyze the insertion of an oxygen atom in a C-C bond (Fraaije et al., 2002). A fourth, less conserved motif common to class B FMOs, is the “FATGY” motif, but can also be referred to as the “ATG-containing”, or “TGY” motif (Stehr et al., 1998).

Functional characterization of different plant FMOs is relatively limited, and predominantly described within model plant species and crops. The best known characterized FMOs include the highly conserved YUCCAs, which perform an oxidative carboxylation of indole-pyruvic acid (IPA) to form the essential hormone auxin (indole acetic acid; IAA; Dai et al., 2013; Mashiguchi et al., 2011; Zhao et al., 2001), and FMO1 which performs a *N-*hydroxylation of pipecolic acid to from the mobile immune signal, *N-*hydroxy pipecolic acid (Hartmann et al., 2018; Holmes et al., 2019). Beyond these well-known FMOs, other characterized members are generally found to perform C-, N- and S- oxidizing reactions within plant specialized metabolism (Florean et al., 2023, 2025; Holmes et al., 2019; Jiang et al., 2025; Kong et al., 2016; Thodberg et al., 2020; Valentino et al., 2020; Won et al., 2011; Chapter IV). Given the diverse and essential catalytic roles these characterized FMOs possess, it is likely that other FMOs carry out key roles in general and specialized metabolism important for plant performance and survival. Further functional analysis of plant FMOs would be greatly aided by a detailed phylogenetic analysis to determine the evolutionary nature and diversity of this important family. Mapping the evolutionary landscape of FMOs will aid in placing newly identified members into clades of shared homology, and can provide clues of shared catalytic functionality. Historically an ad hoc naming system of different FMOs has been employed, and may offer some confusion. For example, FMO1 from *Arabidopsis thaliana* and Garlic (*Allium sativum*) are evolutionary distinct, and catalyze respective N- and S-hydroxylation reactions. Furthermore, these FMO1s and are evolutionary distant to the similarly named mammalian FMO1. At present, phylogenic analysis of plant FMOs contains a small number of plants species, include species biased towards model angiosperms, or has focused on individual FMO families (Carrillo-Carrasco et al., 2023; Jiang et al., 2025; Thodberg et al., 2020, Chapter IV).

The most comprehensive phylogenetic analysis of class B FMOs includes a diverse array of FMOs representing different kingdoms of life (Nicoll & Mascotti, 2023). Six major clades were identified, four of which contained plant-specific FMOs: Clade II which contains the plant S-Ox FMO family, Clade III contains the YUCCAs, Clade IV contains the plant N-Ox family and lastly, the BVMOs (historical name preserved). Although the phylogenetic analysis conducted by Nicoll & Mascotti (2023) provides valuable insights into the evolutionary history of class B FMOs across life, the plant lineage remains underexplored and underrepresented with only three model plants species included in the analysis; a eudicot (*Arabidopsis thaliana*), monocot (*Oryza sativa*) and a bryophyte (*Physcomitrium patens*). A more comprehensive, plant-centric phylogenetic investigation is necessary to uncover lineage-specific adaptations and functional diversification within this enzyme family in the plant kingdom. By expanding the dataset to include a broader range of species and sequences, it is possible to refine our understanding of class B FMO evolution and their roles in plant metabolism. This study therefore aims to address this gap by conducting a detailed phylogenetic analysis of class B FMOs in plants, providing a nuanced perspective on their evolutionary history and functional specialization. Furthermore, we map the structural diversity and placement of FMO-specific motifs and describe their conservation and variation across plant class B FMOs. By refining the definition and structural context of these conserved motifs, this study provides a valuable framework to aid researchers in more accurately identifying class B FMOs, thereby facilitating their phylogenetic classification and functional prediction. Understanding the evolutionary and structural diversification of class B FMOs in plants is crucial for elucidating their roles in specialized metabolic pathways, where monooxygenases frequently catalyze key regio- and stereo-specific oxidation reactions. As central enzymes in hormone biosynthesis, defense compound production, and secondary metabolism, class B FMOs represent an underexplored reservoir of enzymatic functions with potential applications in both plant biology and synthetic biochemistry.

## Results

### A wide net catches many: navigating the phylogenetic blast out of class B FMOs

To ensure a comprehensive and balanced phylogenetic analysis of class B FMOs across embryophyte, the Lifemap server (De Vienne, 2016) was used to select a broad representation of plant species across the plant kingdom (Supplementary Figure 1). A total of 78 green algae and plant species were selected, spanning Chlorophyta, Charophyta, Liverworts, Mosses, Hornworts, Lycophytes, Monilophytes (ferns), Gymnosperms, and a diverse range of Angiosperms (**Figure 2A**). To specifically identify class B FMOs across the plant kingdom, a dedicated pipeline was designed to capture FMO diversity, including potentially undescribed FMO families (Figure 3A). Representative class B FMOs from YUCCAs, BVMO, S-Ox, and N-Ox families across different plant lineages were used as BLASTP queries against collected proteomes and online repositories (Table S1, Table S2), resulting in an initial dataset of 2,339 sequences (E-value <10^−3^). To facilitate reliable manual curation, this dataset was subdivided into smaller, less diverse alignments, as the full set was too divergent to assess pseudogenes, misannotations, and transcript variants effectively. Each subset was then manually inspected and curated, ultimately reducing the dataset to 1,642 high-confidence sequences. A maximum likelihood phylogenetic analysis was then performed on the curated dataset, resulting in the identification of 19 distinct phylogenetic clades (Figure 3B). These phylogenetic groups were first delineated based on tree topology and then annotated according to their best reciprocal BLAST hits (≥50% identity), which included a mixture of characterized enzymes (e.g., YUCCAs, lysine-specific demethylases, glutamate synthase), putative homologs (e.g., monooxygenase 2-like proteins), and several uncharacterized sequences (Figure 3B). Besides the class B FMOs, several of these putatively identified enzyme groups are known to contain FAD and are classified as flavoproteins, including lysine-specific demethylase, glutamate synthase, and protoporphyrinogen oxidase (Brzezowski et al., 2019; Perillo et al., 2020; Van Den Heuvel et al., 2004).

**Figure 2.**
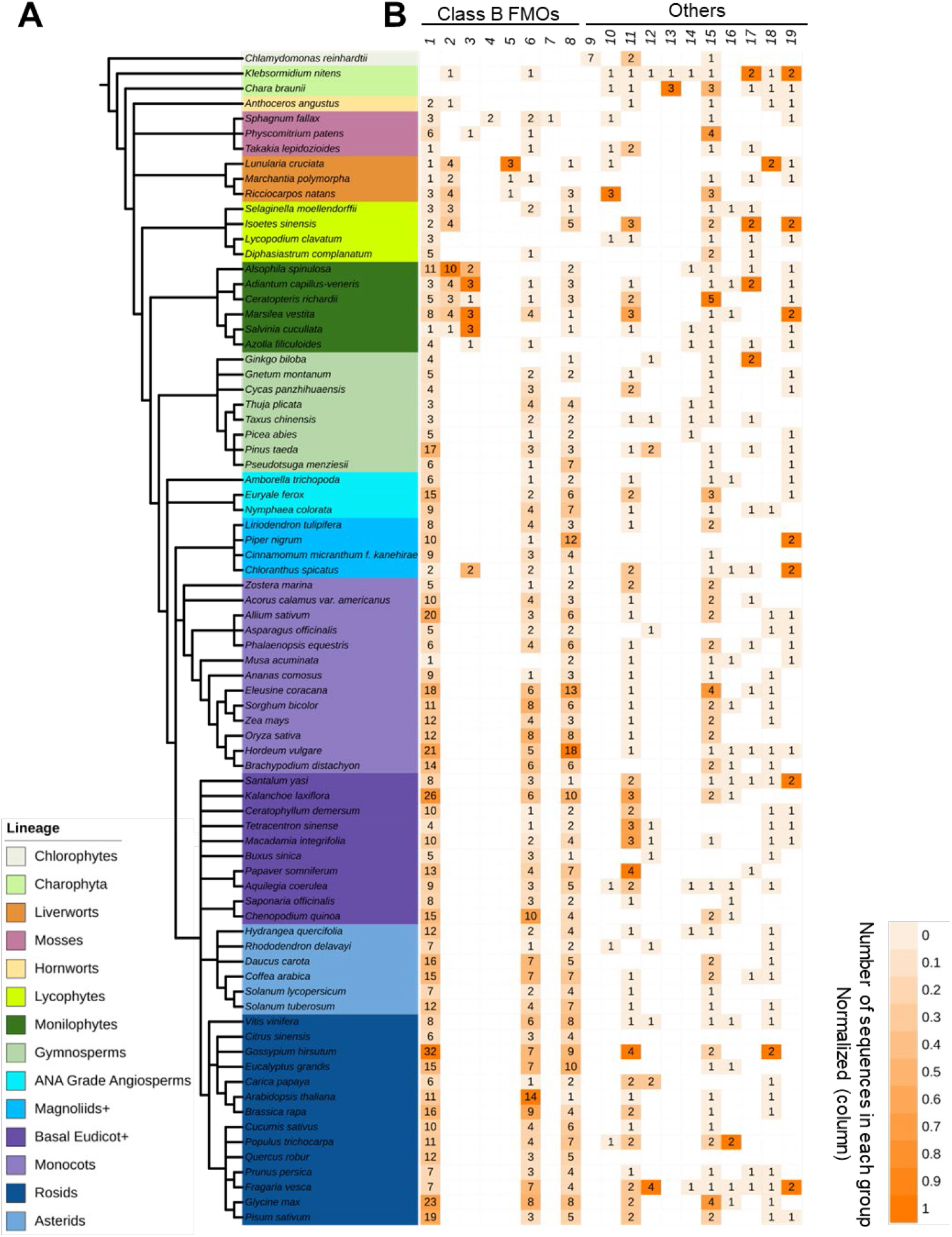
Taxonomic tree of selected species and corresponding sequence distribution across phylogenetic groups. **A)** The taxonomic tree displays the evolutionary relationships among the 78 species included in this study, representing a broad diversity within the plant kingdom. **B)** Heatmap illustrates the distribution of sequences from each species across phylogenetic groups of an enzyme family. Heatmap columns represent phylogenetic groups with values normalized per column to account for differences in total sequence counts. See Figure 3 for corresponding protein family names 1 – 19. Taxonomic overview and sequence distribution can be accessed at: https://itol.embl.de/shared/PlantEcoPhys

**Figure 3.**
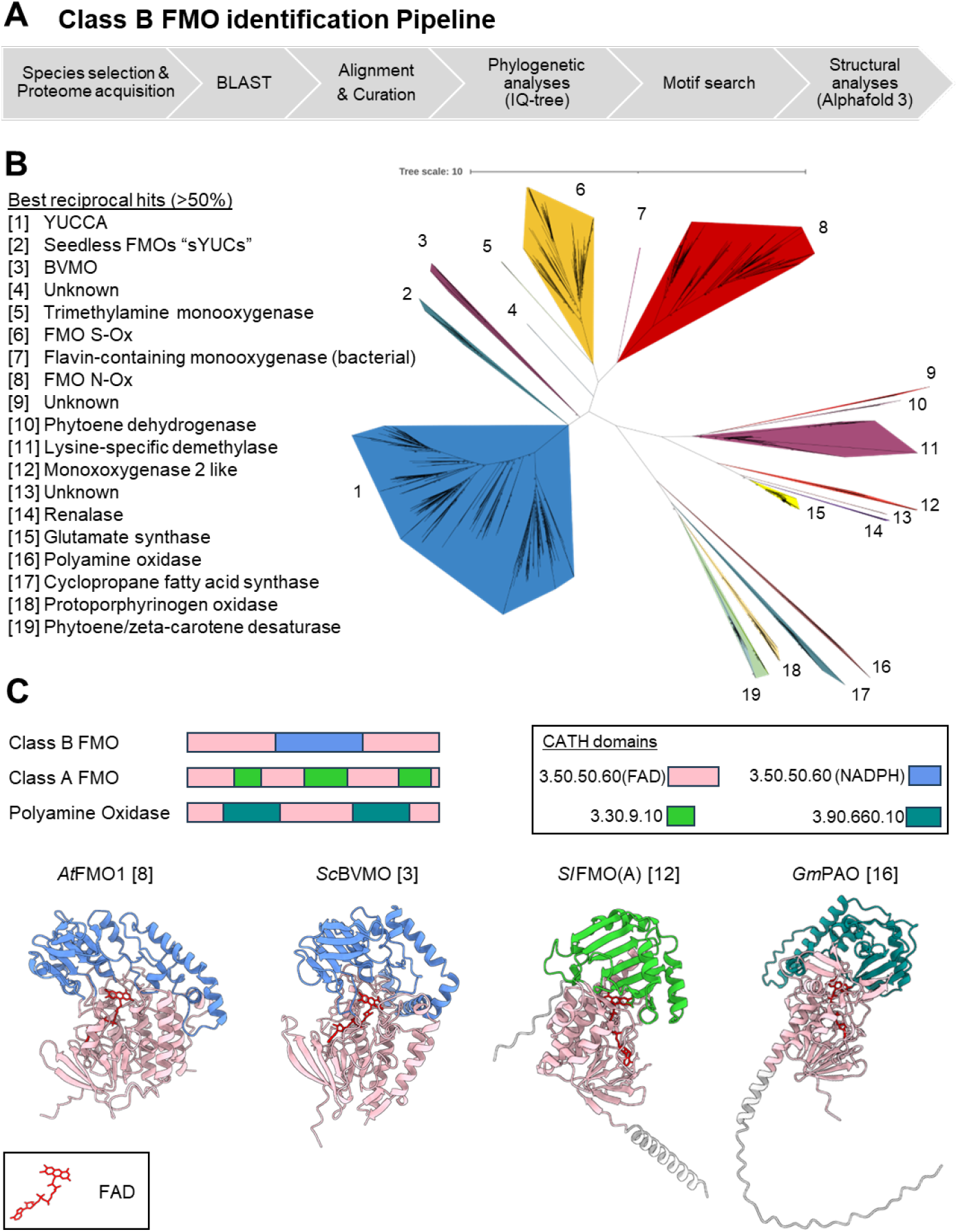
Identification and characterization of Class B FMOs through phylogenetic and structural analysis. **(A)** Pipeline for phylogenetic and structural analysis. The workflow illustrates the process from species selection to sequence analysis, including BLAST searches, multiple sequence alignment (MAFFT), phylogenetic inference (IQ-TREE with ModelFinder (JTT+R10), a motif search and structural analysis of domain architecture. **(B)** Initial phylogenetic tree of sequences acquired by BLAST with class B FMO queries. The tree was constructed using Maximum Likelihood inference in IQ-TREE based on a MAFFT alignment. Nineteen phylogenetic groups were identified, representing evolutionary relationships among FMO sequences. Best reciprocal BLAST hits from NCBI indicate potential enzyme functions. Phylogenetic tree (including bootstrap values) is available at https://itol.embl.de/shared/PlantEcoPhys. **(C)** Structural validation of class B FMO domain architecture. AlphaFold 3 models of four representative enzymes are shown with FAD as a ligand (red) each belonging to distinct phylogenetic clades [Shown in squared brackets]. CATH domains are color-coded: the first Rossmann fold in pink and the second in cornflower blue. CATH domain distinctive of class A FMOs (3.30.9.10) is colored in green whilst 3.90.660.10 from polyamine oxidases is colored in dark cyan

An examination of the resultant phylogenetic tree revealed several emerging patterns. First, the three well-characterized plant class B FMO families (i.e., YUCCAs, FMO N-Ox, and FMO S-Ox) showed broad representation across most plant lineages (Figure 2B, 3B). YUCCAs were identified in all non-algal plant species analyzed, while N-Ox and S-Ox FMOs were generally present in vascular plants and sporadically found in non-vascular groups such as bryophytes and lycophytes. Within these families, angiosperms displayed an expansion in sequence number, suggesting lineage-specific diversification. Notably, the YUCCA and FMO N-Ox clades lacked sequences from algal lineages (Figure 2B). Furthermore, no *Chlamydomonas reinhardtii* sequences were identified within the first eight phylogenetic groups classified as class B FMOs, suggesting either a true absence or significant divergence of these enzymes in this algal species. Beyond these canonical FMO families, several clades with the best reciprocal BLAST hits to non-class B FMO enzymes also exhibited broad phylogenetic distribution. Notably, clades 11 and 15, corresponding to lysine-specific demethylases and glutamate synthases, respectively, were represented across nearly all major plant lineages; from chlorophytes to angiosperms (Figure 2B). In contrast, other clades such as clade 12 (monooxygenase 2-like) and clade 14 (renalase) showed more limited representation, appearing only in a restricted subset of taxa.

To assess if any of the retrieved sequences represented novel class B FMO families, an unbiased analysis of all 19 phylogenetic groups was conducted to assess the prevalence of canonical class B FMO motifs. Sequences were selected for further investigation based on the presence of at least two Rossmann fold motifs (GxGxxG/A) and a conserved FMO-identifying motif, characterized by a highly conserved histidine residue. Structural analysis of AlphaFold 3 (Abramson et al., 2024) predicted models for representative sequences revealed that eight phylogenetic groups exhibited the distinctive class B FMO architecture, consisting of two FAD/NAD(P)-binding domains (CATH 3.50.50.60), with the NADPH-binding domain splitting the FAD-binding domain (Figure 2C). Interestingly, phylogenetic groups 12 and 16 (Figure 2B) shared sequence similarities with class B FMOs, possessing two Rossmann fold motifs (GxGxxG/A) and an FMO-identifying motif (FXGXXXHXXXY/F). However, structural comparison using the Foldseek server (Van Kempen et al., 2024) classified representatives from these groups—*Sl*FMO(A) (group 12; XP_069144694.1) and *Gm*PAO (group 16; Glyma.15G276500)—as a class A FMO and a polyamine oxidase, respectively. These findings were further validated through domain architecture analysis using the CATH server (Greene et al., 2007)(Figure 3B). *Sl*FMO(A) from tomato was identified as a class A FMO and appeared in the initial phylogenetic analysis within clade 12, which had monooxygenase 2-like proteins as its best reciprocal BLAST hit (Figure 3B). This family is not broadly represented across plant lineages, with 18 sequences retrieved from 12 taxonomically diverse species (Figure 2B).

### Plant-centric phylogenetic analysis of Class B FMOs

Based on the first navigational phylogenetic output and broad structural comparisons, a second phylogenetic analysis was performed using only the determined plant class B members. This analysis shows that plant class B FMOs are polyphyletic, with eight distinct phylogenetic clades supported by bootstrap values of 100. As expected, the large canonical YUCCA, N-Ox and S-Ox families form major branches in the tree. The other well recognized BVMO family previously represented by only a single characterized member from the moss *P. patens* (Beneventi et al., 2013) was expanded to include new BVMO members from the angiosperm species *Chlorantus spicatus*, and 13 monilophyte members. In addition these, four new FMO families are formally recognized, with relatively narrow representation across the plant kingdom (Figure 2B). One clade was previously suggested to be a sister group to the YUCCA family (Carrillo-Carrasco et al., 2023). This family consists of FMO members from non-seed plants (algae, liverworts, hornworts, lycophytes and monilophytes)(Figure 2B), so we therefore suggest naming this family “Seedless FMOs” [3]. Of the three remaining novel phylogenetic FMO families, two consist exclusively of members from the moss *Sphagnum fallax* (S-OX-like2 [4] and N-Ox-like [7]), and the third family only consists of members from liverwort species *Marchantia polymorpha, Ricciocarpos natans* and *Lunularia cruciata* (S-Ox-like 1 [5]).

### Sequence diversity across key plant Class B FMO motifs

To investigate the structural diversity within class B FMOs, eight representative enzymes were selected, each corresponding to a previously identified family (Figure 4). Structural models were predicted using AlphaFold 3 with FAD as a ligand. Consistent with prior characterizations, all class B FMOs adopt a two-domain architecture, comprising an FAD-binding domain and an NADPH-binding domain. The two dinucleotide-binding domains (CATH 3.50.50.60) were highlighted in the structural models, with the FAD-binding domain colored pink and the NADPH-binding domain in cornflower blue (Figure 5). Class B FMOs are typically characterized by several conserved motifs, including the FAD-binding motif (GxGxxG), the NADPH-binding motif (GxGxxG/A), the FMO-identifying motif (FxGxxxHxxxY/W), and the F/LATGY (or ATG) motif. These motifs were identified in AtYUC10 and highlighted in orange (FAD/NADPH-binding domain), orchid/purple (FMO-ID motif), and aquamarine (F/LATGY motif). However, several of the other representative FMOs did not display these motifs with the expected sequence residues. To systematically assign these motifs across all phylogenetic clades, AtYUC10 was structurally aligned to the other representative structures. This alignment revealed that the structural folds corresponding to the reported conserved motifs were well-preserved across all analyzed FMOs, even when sequence conservation was not immediately evident (Figure S2). Using these superimposed structures, the conserved fold regions were mapped onto each structure, allowing for a structurally informed assignment of motifs across the phylogenetic groups.

**Figure 4:**
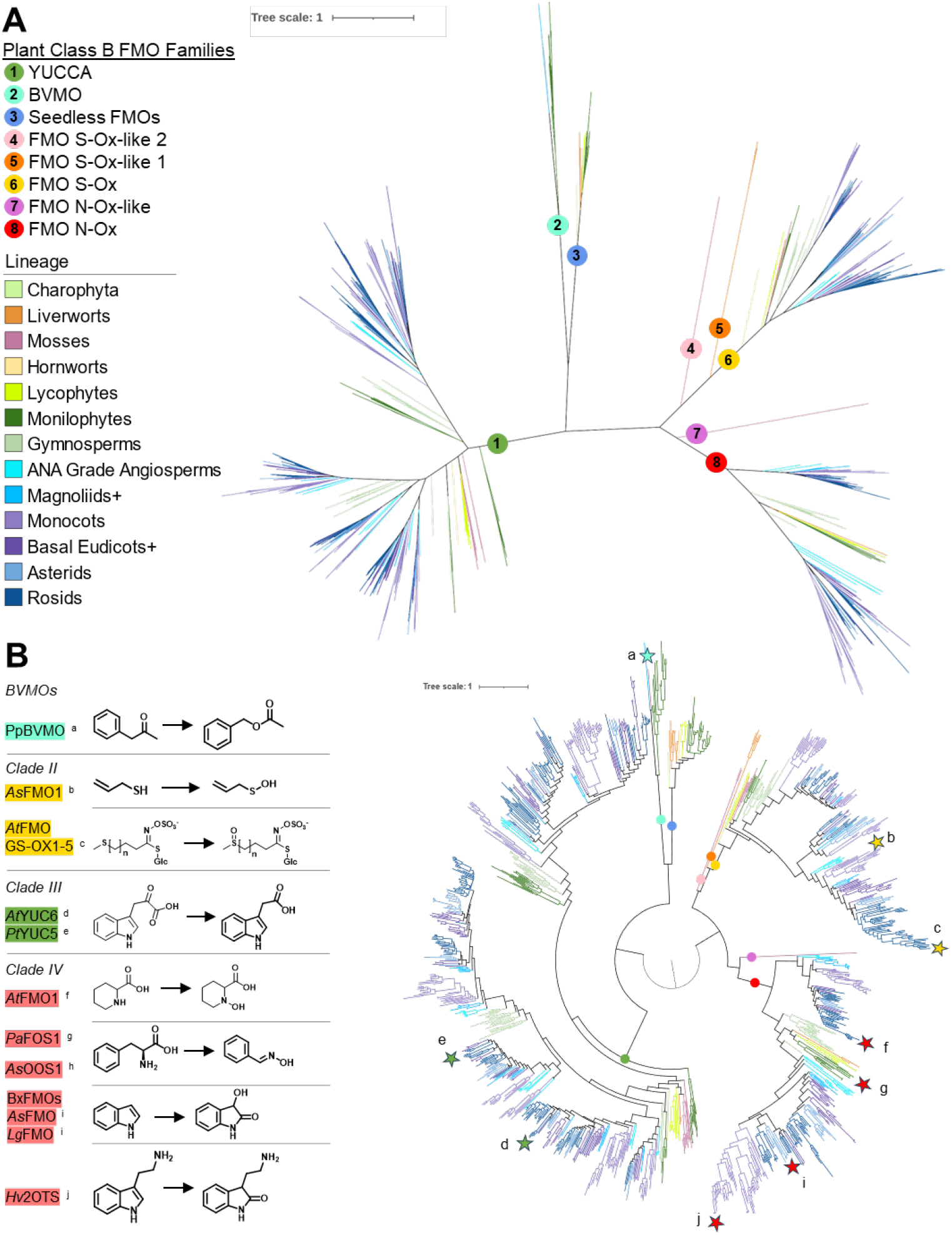
Plant-centric class B FMO phylogeny. **A)** Unrooted maximum likelihood phylogenetic tree identified class B FMOs with eight distinct phylogenetic families denoted by numbered colored circles. **B)** Radial visualization of the phylogenetic analyses rooted at the midpoint, with characterized plant class B FMOs highlighted with letters a-j, and their corresponding catalytic functions listed on the left. Online version of the tree is available at: https://itol.embl.de/shared/PlantEcoPhys

**Figure 5:**
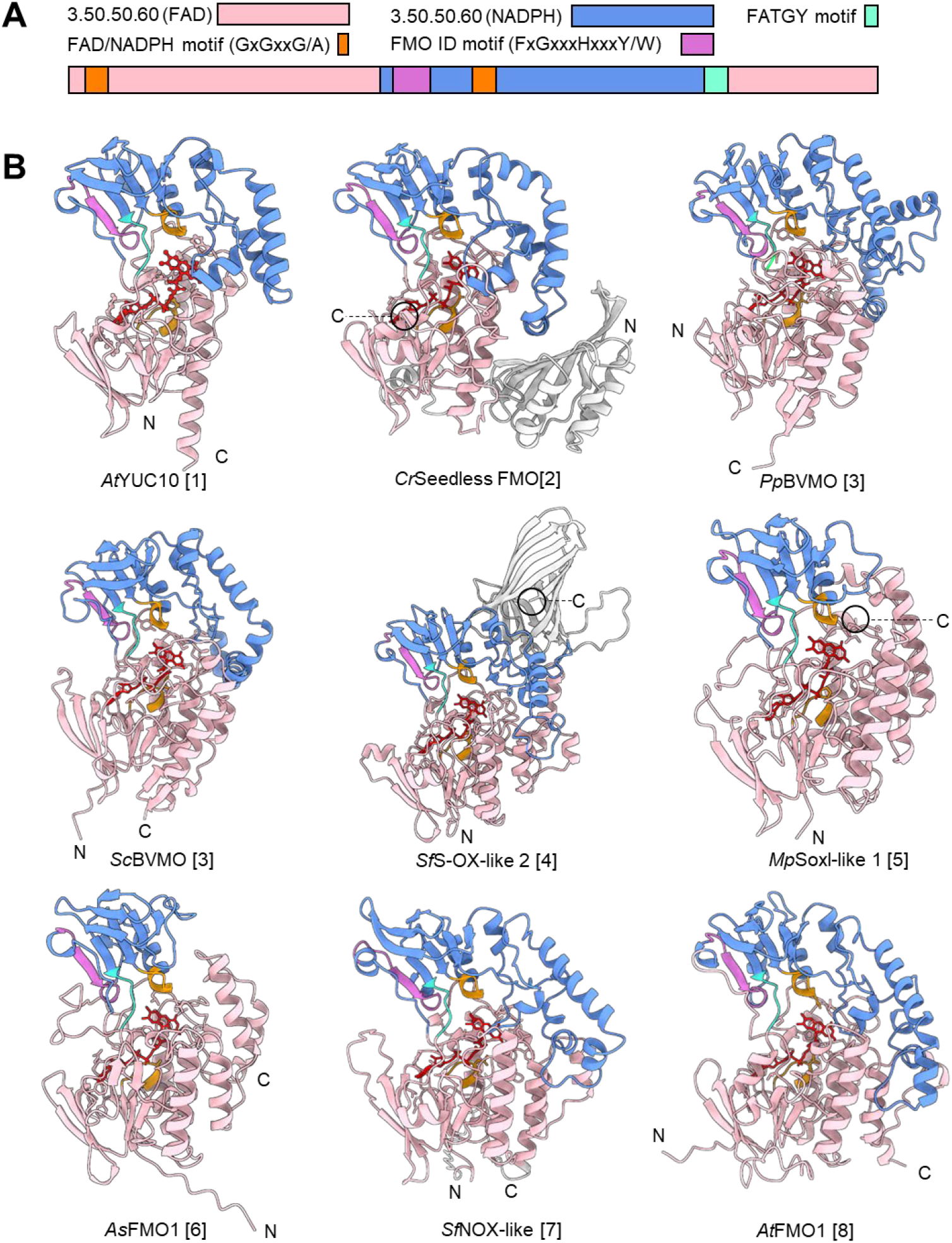
Structures of identified Class B FMOs. **A)** Class B FMO Schematic of characteristic dinucleotide domain architecture and canonical motifs/folds. **B)** AlphaFold 3 predicted structures of representatives from eight identified phylogenetic families of class B FMOs. The FAD- and NADPH-binding domains are colored in pink and cornflower blue respectively. Structural folds belonging to canonical class B FMO motifs are colored in orange (GxGxxG/A), purple (FMO ID motif; (FxGxxxHxxxY/W) and aquamarine (FATGY). FAD molecule is colored red.

The most pronounced structural differences were observed in the NADPH-binding domain, particularly in the region between the NADPH-binding motif and the FATGY motif, where the size and structural embellishments varied significantly. Notably, despite these variations, the FATGY motif was consistently positioned in the same overall location within the protein, effectively facilitating the transition from the NADPH-binding domain to the second part of the FAD-binding domain. This conserved positioning suggests a structural role in maintaining domain connectivity and functional coordination within class B FMOs. SOx and SOx-like 1 representatives exhibited the simplest NADPH-binding domain, whereas other groups displayed additional α-helical insertions. Notably, the Seedless FMO group possesses a conserved N-terminal domain in addition to the distinct class B FMO domain architecture, denoted as a catalytically inactive SnoaL-like fold (Chánique et al., 2023). Additionally, both SOx-like 2 FMOs from *Sphagnum fallax* are distinguished by a C-terminal β-barrel.

To better understand the evolutionary relationships among the identified FMO families, we performed structural alignments of representative AlphaFold3-predicted models (Figure 1). The Seedless FMO *Ceratopteris richardii* (Ceric.34g041100) was used as a test case to evaluate similarity to known plant FMO families. Structural comparison showed it was most similar to *At*YUC10 (YUCCA) and *At*FMO1 (FMO N-Ox), with RMSD values of 5.3 and 6.8, respectively, while showing lower similarity to the BVMO representative (*Pp*BVMO; RMSD = 9.3) (Table S3). These values indicate that while the Seedless FMO shares general FMO fold characteristics, the family does not exhibit greater structural similarity to YUCCAs compared with other class B FMOs, supporting its position as a distinct family.

Structural modeling confirmed that the overall folds associated with key class B FMO motifs are conserved across all plant FMO families, providing a stable framework for comparing sequence-level divergence and motif variation. The canonical FAD-binding Rossmann fold (GxGxxG) was consistently conserved across all families. In contrast, the NADPH-binding motif displayed greater variability. While YUCCAs and BVMOs retained the central glycine, most other families showed a GxxxSA or GxxxSG variant with a conserved serine residue. The FMO-identifying motif (FxGxxxHxxxY) also exhibited family-specific substitutions, such as a tryptophan (W) replacing phenylalanine (F) in the S-Ox family (WxGxxxHxxxY), and a loss of the central histidine was observed in S-Ox-like 2 and N-Ox-like families. The FATGY motif, considered less conserved, showed a more consistent pattern with a conserved (A/C)TG core: ATG in YUCCAs, BVMOs, FMO N-Ox and Seedless FMOs, and CTG in S-Ox and S-Ox-like families (Table 1).

**Table 1.**
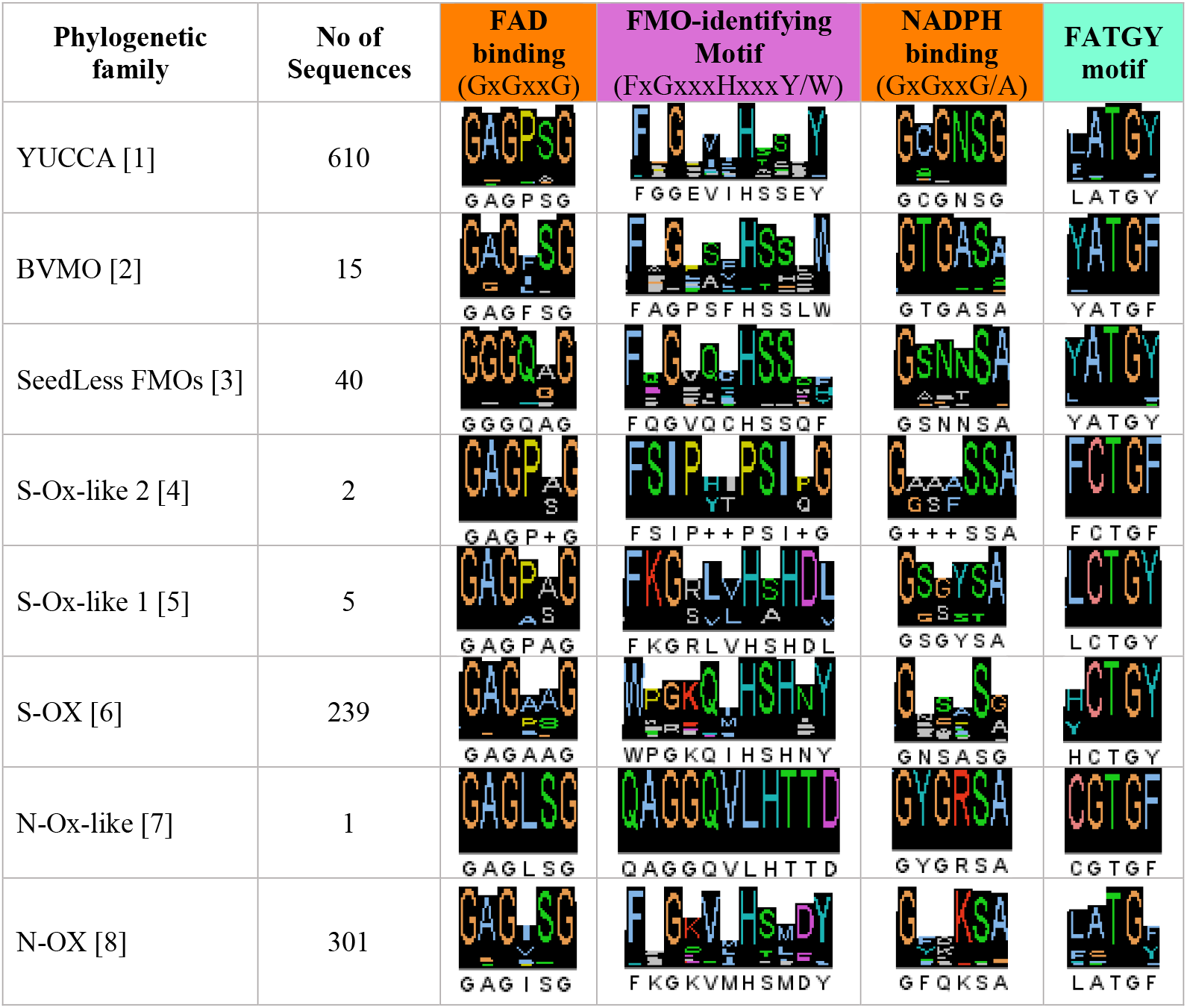
Revaluation conserved Plant Class B FMO motifs represented by logos of different families.

## Discussion

In this study, we present a comprehensive phylogenetic and structural analysis of class B flavin-containing monooxygenases (FMOs) across the plant kingdom. To place these enzymes in a broader evolutionary context, we also examine the diversity of class B FMOs alongside other flavin-dependent proteins that share partial sequence similarity. A large-scale BLAST search using known plant class B FMO sequences returned a wide range of flavin-related proteins, many of which shared conserved regions corresponding to the FAD-binding domain. For example, proteins from the Polyamine oxidase and class A FMO clades (Figure 3C) both possess two Rossmann folds and an FMO-like identifying motif, and share approximately 30– 50% sequence identity across 30–60 aligned amino acids relative to class B FMO queries. Similarly, the glutamate synthase family was captured due to common features in the FAD-binding region, with ∼40–50% sequence identity over alignment lengths of 40–70 amino acids. These hits reflect general sequence conservation involved in FAD binding rather than evidence of a shared evolutionary origin with class B FMOs. Importantly, this analysis establishes a clear framework to distinguish class B FMOs from other flavin-containing protein families in plants.

Interestingly, the tomato class A FMO identified in this study (*Sl*FMO(A); Figure 3C) is distinct from the previously reported tomato class A FMO, *Sl*TNH1, described by Liscombe et al., (2022). *Sl*FMO(A) has been shown to negatively regulate drought tolerance in tomato through modulation of reactive oxygen species (ROS) homeostasis (Wang et al., 2023). In the present study, this FMO belongs to a clade comprising 18 sequences from 12 species, suggesting the presence of additional class A FMOs across the plant kingdom. It is noted, however, that as the BLAST out did not specifically target class A FMOs, nor did it retrieve the characterized *Sl*TNH1. It is therefore highly likely that class A FMOs have a much wider representation in plants than what is reported here. While *Sl*FMO(A) and *Sl*TNH1 share only 37% sequence identity, structural alignment of their AlphaFold-predicted models revealed a high degree of similarity, with an RMSD of 0.930 Å across 295 atom pairs. This structural conservation despite low sequence similarity highlights the underlying diversity within the class A FMO family and underscores the need for further research to map this family across plant lineages.

Using the genomes of 78 different plant and algae species, eight distinct class B FMO families were identified, including three previously unrecognized clades composed of sequences from non-seed plant species. In addition, the plant BVMO family, previously represented by a single characterized member from moss (Beneventi et al., 2013), has been expanded to include BVMOs from ferns (monilophytes) and a single angiosperm. Furthermore, the phylogenetic placement of an FMO family previously proposed to lie sister to the YUCCA family has been clarified and hereby named SeedLess FMOs. Of the eight class B FMO families identified in this study, only four contain members with previously characterized enzymatic functions. These families map onto clades II, III, and IV of the class B FMO phylogeny reported by Nicoll and Mascotti (2023). Clade II includes the S-oxygenating FMOs such as *As*FMO1 and *At*FMO GS-OX1–7, and likely also includes the newly identified S-Ox-like 1 and 2 families. Clade III contains the YUCCA family, while Clade IV is composed almost entirely of plant sequences and includes the catalytically diverse FMO N-Ox family, with the FMO N-Ox-like group also likely belonging to this clade (Nicoll & Mascotti, 2023, Figure 4B).

Some class B FMO families, namely the YUCCA, N-OX and S-OX families, are highly prevalent and diverse across the plant kingdom, particularly within angiosperms (Figure 2B). The YUCCA family is especially large, with many YUCCA genes retained following many whole gene duplication events (see Chapter V for more detail). All characterized YUCCAs from different plant species catalyze a conserved reaction: the oxidative decarboxylation of IPA to form IAA, an essential hormone (Figure 4B). Accordingly, the high copy number of YUCCAs is largely attributed to the fine-tuned regulation of auxin across plant development and in response to a wide range of environmental factors (Chapter V). Some YUCCAs have also been reported to possess thiol reductase activity which aids the mediation of reactive oxygen species upon environmental (Cha et al., 2015; Wang et al., 2022), but it is currently unknown how conserved this role is across the YUCCA family.

In contrast to YUCCAs, the high copy number of N-OX and S-OXs in plants may be due to species-specific adaptations and unique responses to environmental factors. Characterized N-OX family members perform *N-* or *C-*hydroxylations, with their catalytic products important for plant defense and immunity. *At*FMO1 is as a key regulator of systemic acquired resistance by catalyzing the *N-*hydroxylation of pipecolic acid to produce *N-*hydroxy-pipecolic acid (NHP), a crucial immune signaling molecule (Hartmann et al., 2018; Holmes et al., 2019). FOS1 from the fern species *Phlebodium aureum* and *Pteridium aquilinum*, and OOS1 from Darwin’s orchid (*Phalaenopsis equestris*) catalyze successive *N-*hydroxylations of an amino acid to form its corresponding aldoxime. Oxime synthase functionality is case of convergent evolution whereby the trait has evolved independently in ferns and in the orchid, and for different purposes (Sánchez-Pérez & Neilson, 2024). In ferns the oxime is formed as an intermediate for cyanogenic glycoside production (Thodberg et al., 2020), while the oximes are emitted from Darwin’s orchid to attract a specific pollinating moth (Jiang et al., 2025); chapter VI). N-Ox *C-*hydroxylation capacity is represented by characterized FMOs from different eudicot species involved in the biosynthesis of benzoxazinoids. These FMOs catalyze two sequential hydroxylations of indole to form hydroxy indolin-2-one (Florean et al., 2023, 2025). In a similar catalytic role, a recently characterized FMO from barley (*Hordeum vulgare*) hydroxylates tryptamine to produce 2-oxo-tryptamine and produced in response to pathogen infection (Chapter III). Fewer S-Ox FMOs have been characterized, but all are involved in sulfur oxidation. *At*GS-OXs are involved in glucosinolate biosynthesis by catalyzing the oxygenation of sulfur atoms (Hansen et al., 2007; Kong et al., 2016) while *As*FMO1 catalyzes a sulfur oxidation in the biosynthesis of allicin (Valentino et al., 2020).

In contrast to the large YUCCA, N-Ox and S-Ox families, the other families show a highly restricted distribution (Figure 2B). Notably, the novel class B FMO families FMO N-Ox-like [7] and FMO S-Ox-like 2 [4] are comprised exclusively of sequences from the moss *S. fallax*. This highly restricted taxonomic distribution raises intriguing questions about their evolutionary origins. One potential explanation lies in the unique genome architecture of the *Sphagnum* genus; where recent comparative studies have shown that *Sphagnum* genomes exhibit no gene collinearity with any other reference genome to date (Healey et al., 2023). This suggests that *Sphagnum* genomic architecture may reflect a more ancient and independently evolving lineage with unique FMO representatives. The absence of collinearity could be the result of retained ancestral genome structures or extensive rearrangements accumulated over deep evolutionary time. An alternative hypothesis is that these gene families arose via horizontal gene transfer (HGT) from microbial sources. Reciprocal BLAST analyses for the N-Ox-like family returned highest similarity scores to bacterial sequences, including *Geodermatophilaceae bacterium* (56.32% identity) and *Actinomycetospora soli* (53.28%), supporting a potential bacterial origin. In contrast, the S-Ox-like 2 family yielded its top BLAST hit to a fungal sequence from *Botryobasidium botryosum* (42.42%), raising the possibility of a distinct fungal contribution. This interpretation is further supported by phylogenetic context: *Sphagnum* sequences are found within the broader moss clades of the S-Ox and YUCCA families, whereas no moss species possess members of the N-Ox family. The prevalence of BVMOs in plants may also be attributed to HGT given that the members of this family are present in distinct but restricted plant lineages of moss, monilophytes and a single angiosperm. A reciprocal BLAST using the BVMO sequence from *Physcomitrium patens* returned ∼64% identity to a homolog from the moss *Ceratodon purpureus* indicating that BVMOs are present in additional moss species beyond *P. patens*. Interestingly, the *Pp*BVMO has ∼62% identity to a bacterial sequence from *Blastococcus deserti* supporting a bacterial origin. Similarly, a BVMO sequence from the monilophyte *Salvinia cucullata* yielded top hits to a bacterial sequence from an *Acidobacteriota* species (∼45% identity). In contrast, the angiosperm BVMO from *Chloranthus spicatus* showed the highest similarity (∼78% identity) to a fungal sequence from *Zymoseptoria tritici*. These findings support the hypothesis that BVMO genes were acquired independently in different plant lineages through multiple horizontal gene transfer events from diverse microbial sources. As the large plant YUCCA family is also widely accepted to arise from HGT (Bowman et al., 2021, Chapter V), this phenomenon has been an important evolutionary innovation for class FMO recruitment within plant metabolism.

The phylogenetic analysis of plant-specific class B FMOs presented here has enabled the reclassification of a lesser-known family previously referred to as “sister YUCCAs”, or “sYUCs”. This family was initially identified as putative YUCCA-like enzymes, and proposed to position evolutionary to the important auxin-biosynthesizing YUCCA family (Carrillo-Carrasco et al., 2023). Phylogenetic and structural analyses, however, clearly demonstrate that this family is an evolutionarily independent lineage within the class B FMO superfamily, and we therefore designate it a new name of “SeedLess FMOs” as it apparently lacks related FMO members from Gymnosperms and Angiosperms. To date, no members of the SeedLess FMO family have been functionally characterized. Their original identification was based on transcriptomic data from the 1KP project (One Thousand Plant Transcriptomes Initiative, 2019) providing strong evidence that these genes are actively transcribed across a wide range of non-seed plant species. This widespread expression implies that SeedLess FMOs may carry out conserved and potentially important biological roles in early-diverging plant lineages. To investigate their expression dynamics in more detail, we analyzed the two SeedLess FMO genes from *Marchantia polymorpha* (Mapoly0122s0026 & Mapoly0052s0062) using the BAR eFP Browser. Both genes are broadly co-expressed across a variety of tissues, with notably higher expression levels in the archegoniophore and antheridiophore, the female and male reproductive structures, respectively. Furthermore, both genes show clear induction in response to abiotic stresses, with particularly strong upregulation of Mapoly0052s0062 under specific environmental conditions (Tan et al., 2023, Figure S3). These patterns suggest potential roles in reproduction and stress adaptation, although functional studies will be required to validate these hypotheses. Interestingly, a FoldSeek search (Van Kempen et al., 2024) of a representative seedless FMO from *C. richardii* against crystallized proteins, identified an FMO from the bacteria *Janthinobacterium svalbardensis* (*Js*FMO), which contains the same SnoaL-like N-terminal fold common to the SeedLess FMO family. *Js*FMO possesses the ability to perform Baeyer-Villiger (BV) oxidations (Chánique et al., 2023) and given the structural similarity, raises the intriguing possibility that SeedLess FMOs may share a comparable catalytic mechanism. While functional data are currently lacking, the closest phylogenetic neighbors to SeedLess FMOs – the BVMOs and YUCCAs – are also associated with BV-type oxidations. Taken together, these observations suggest that SeedLess FMOs could represent an additional, uncharacterized branch of BV-oxidizing enzymes in early-diverging land plants, warranting further biochemical investigation.

The structural analyses performed in this study showed that the overall domain architecture is conserved across class B FMOs, particularly in the core orientation of the FAD- and NADPH-binding domains, though notable structural variability was observed within the NADPH-binding domain (Figure 5). The conserved elements are likely essential for maintaining the fundamental catalytic function of these enzymes, whereby the bound FAD is reduced by NADPH and reacts with dioxygen to form a moderately stable C4a-(hydro)peroxy FAD intermediate. This intermediate reacts with the substrate to perform the catalytic reaction. In the absence of a substrate the C4a-(hydro)peroxy intermediate decays into oxidized FAD and hydrogen peroxide; a process known as uncoupling (Ziegler, 1988). In contrast, the more structurally diverse regions may contribute to the functional diversification observed across different FMO families. This variable portion could influence substrate specificity by acting as a gatekeeping module that regulates access to the active site. This variable region may also be involved in determining the oligomeric state of the protein. Most purified or structurally characterized class B FMOs has been shown to form homodimers (Alfieri et al., 2008; Cho et al., 2011; Eswaramoorthy et al., 2006; Thodberg et al., 2020). One exception is the S-Ox FMO – *As*FMO1 – from garlic (*Allium sativum). As*FMO1 is the only plant-derived class B FMO for which a crystal structure has been reported, and both crystallographic data and size exclusion chromatography confirmed that it forms a monomer in solution (Valentino et al., 2020). Interestingly, *As*FMO1 also possesses the smallest and least elaborated NADPH-binding domain among the predicted structures and may suggest a potential link between structural truncation and monomeric behavior. Taken together, these findings raise the possibility that the structurally variable portion of the NADPH domain may influence dimerization or mediate protein-protein interactions. To date, very few biochemical studies on plant FMOs have been reported. Aside from the aforementioned *As*FMO1, only *At*YUC6 has been biochemically characterized in any detail (Dai et al., 2013). This lack is likely due to issues relating to FMO expression and purification, whereby FMOs have been referred to as being difficult to handle (Thodberg & Jakobsen Neilson, 2020). Nonetheless, biochemical studies focusing on the dimeric interfaces and overall quaternary structure of plant class B FMOs will be necessary to determine whether structural variation in different regions contributes to functional specialization.

## Conclusions

This study presents a comprehensive phylogenetic and structural analysis of class B FMOs in plants, identifying eight distinct families, including three newly recognized groups found exclusively in non-seed plants. Structural modeling showed that all plant class B FMOs share a conserved domain architecture composed of two dinucleotide-binding domains. However, considerable structural diversification was observed across families, particularly within the NADPH-binding domain. In addition, we define a refined set of conserved FMO-specific motifs by anchoring the analysis in structurally conserved folds rather than amino acid sequence alone. These motifs offer a tool for phylogenetic annotation and discovery of uncharacterized plant class B FMOs in future studies. Altogether, the expanded phylogenetic analyses and insights into structural variation provide a foundation for more precise characterization of class B FMOs in plants.

## Methods

### Dataset Assembly

A total of 78 species were selected to ensure broad and comprehensive representation across the Streptophyta lineage (Figure S1, Table S2), with a prerequisite of a fully sequenced genome. Genomes were retrieved from publicly available repositories, including Phytozome (Goodstein et al., 2012), PlantGenie (Sundell et al., 2015), Phycocosm (Grigoriev et al., 2021), and FernBase (Li et al., 2018). If a genome was unavailable in these databases, predicted peptide FASTA files were obtained from other sources (Table S1). Representative sequences from each class B FMO family were selected, including those from key model organisms: *Marchantia polymorpha* (liverwort), *Physcomitrium patens* (moss*), Selaginella moellendorffii* (lycophyte), <u>*Ceratopteris richardii*</u> (monilophyte), *Picea abies* (gymnosperm), *Arabidopsis thaliana* (dicot), and *Hordeum vulgare* (monocot) (Table S2). These sequences were used as BLAST queries at the amino acid level against each of the 78 genomes individually. Local BLAST searches were performed against retrieved proteomes, and hits with an e-value ≤ 10^−3^ were retained. Sequences were aligned using MAFFT v7 (Katoh & Standley, 2013), and alignments were refined by separating less diverse sequences into distinct alignments. Manual curation was conducted to remove misannotated proteins and pseudogenes exhibiting large insertions, deletions, or frameshift mutations.

### Motif analysis

The identification of canonical class B FMO motifs across phylogenetic groups 1–19 was conducted using the motif search tool in CLC Main Workbench 20.0.4 (ref). The motifs used included the FAD-binding motif (GxGxxG), NADPH-binding motif (Gx(x)GxxG/A), FMO-identifying motif (FxGxxxHxxxY/F), BVMO-identifying motif (FxGxxxHxxxW), and FATGY motif (F/L/YATGY), all applied with an 80% accuracy threshold. To generate motif logos for class B FMO families (groups 1–8), multiple sequence alignments were performed using MAFFT and visualized in Jalview. Sequences lacking information at the motif sites were excluded from conservation analyses.

### Phylogenetic analysis

Phylogenetic relationships were inferred using maximum-likelihood (ML) analysis based on multiple sequence alignments (MSA) generated with MAFFT v7. The initial phylogenetic analysis included 1,642 sequences, while the class B FMO phylogenetic analysis incorporated 1,273 sequences. ML phylogenetic inference was conducted using IQ-TREE with ModelFinder selecting the optimal substitution models: JTT+R10 for the initial analysis and JTT+F+R10 for the class B FMO analysis. All phylogenetic trees were inferred with 1,000 bootstrap replicates to assess branch support.

### Structural analysis

All structural models were predicted using AlphaFold 3 (Abramson et al., 2024), incorporating FAD as a ligand. Structural analyses and visualizations were conducted with ChimeraX 1.9 (Meng et al., 2023). To supplement manual domain architecture determination, the CATH domain server (Greene et al., 2007; Orengo et al., 1997) was used. Accession of predicted structures (AtYUC10[1]; AT1G48910, CrSeedless FMO[2]; Ceric.34G041100, PpBVMO[3]; Pp6c18_10300, ScBVMO[3]; Sacu_v1.1_s0081.g018001, SfS-OX-like 2[4]; Sphfalx02G048800, MpS-Ox-like 1[5]; Mapoly0020s0006 AsFMO1[6]; Asa8G05386.1, SfN-Ox-like[7]; Sphfalx14G087800, AtFMO1[8]; AT1G19250). Structural alignments and root mean square deviation (RMSD) calculations (Table Sx) were performed using the Matchmaker tool (Meng et al., 2006) in ChimeraX, with no pruning of long atom pairs.

## References

Abramson, J., Adler, J., Dunger, J., Evans, R., Green, T., Pritzel, A., Ronneberger, O., Willmore, L., Ballard, A. J., Bambrick, J., Bodenstein, S. W., Evans, D. A., Hung, C.-C., O’Neill, M., Reiman, D., Tunyasuvunakool, K., Wu, Z., Žemgulytė, A., Arvaniti, E., … Jumper, J. M. (2024). Accurate structure prediction of biomolecular interactions with AlphaFold 3. Nature, 630(8016), 493–500. 10.1038/s41586-024-07487-w

Alfieri, A., Malito, E., Orru, R., Fraaije, M. W., & Mattevi, A. (2008). Revealing the moonlighting role of NADP in the structure of a flavin-containing monooxygenase. Proceedings of the National Academy of Sciences, 105(18), 6572–6577. 10.1073/pnas.0800859105

Beneventi, E., Niero, M., Motterle, R., Fraaije, M., & Bergantino, E. (2013). Discovery of Baeyer– Villiger monooxygenases from photosynthetic eukaryotes. Journal of Molecular Catalysis B: Enzymatic, 98, 145–154. 10.1016/j.molcatb.2013.10.006

Bowman, J. L., Flores Sandoval, E., & Kato, H. (2021). On the Evolutionary Origins of Land Plant Auxin Biology. Cold Spring Harbor Perspectives in Biology, 13(6), a040048. 10.1101/cshperspect.a040048

Brzezowski, P., Ksas, B., Havaux, M., Grimm, B., Chazaux, M., Peltier, G., Johnson, X., & Alric, J. (2019). The function of PROTOPORPHYRINOGEN IX OXIDASE in chlorophyll biosynthesis requires oxidised plastoquinone in Chlamydomonas reinhardtii. Communications Biology, 2(1), 159. 10.1038/s42003-019-0395-5

Carrillo-Carrasco, V. P., Hernandez-Garcia, J., Mutte, S. K., & Weijers, D. (2023). The birth of a giant: Evolutionary insights into the origin of auxin responses in plants. The EMBO Journal, 42(6), e113018. 10.15252/embj.2022113018

Cha, J.-Y., Kim, W.-Y., Kang, S. B., Kim, J. I., Baek, D., Jung, I. J., Kim, M. R., Li, N., Kim, H.-J., Nakajima, M., Asami, T., Sabir, J. S. M., Park, H. C., Lee, S. Y., Bohnert, H. J., Bressan, R. A., Pardo, J. M., & Yun, D.-J. (2015). A novel thiol-reductase activity of Arabidopsis YUC6 confers drought tolerance independently of auxin biosynthesis. Nature Communications, 6(1), 8041. 10.1038/ncomms9041

Chánique, A. M., Polidori, N., Sovic, L., Kracher, D., Assil-Companioni, L., Galuska, P., Parra, L. P., Gruber, K., & Kourist, R. (2023). A Cold-Active Flavin-Dependent Monooxygenase from Janthinobacterium svalbardensis Unlocks Applications of Baeyer–Villiger Monooxygenases at Low Temperature. ACS Catalysis, 13(6), 3549–3562. 10.1021/acscatal.2c05160

Cho, H. J., Cho, H. Y., Kim, K. J., Kim, M. H., Kim, S. W., & Kang, B. S. (2011). Structural and functional analysis of bacterial flavin-containing monooxygenase reveals its ping-pong-type reaction mechanism. Journal of Structural Biology, 175(1), 39–48. 10.1016/j.jsb.2011.04.007

Dai, X., Mashiguchi, K., Chen, Q., Kasahara, H., Kamiya, Y., Ojha, S., DuBois, J., Ballou, D., & Zhao, Y. (2013). The Biochemical Mechanism of Auxin Biosynthesis by an Arabidopsis YUCCA Flavin-containing Monooxygenase. Journal of Biological Chemistry, 288(3), 1448–1457. 10.1074/jbc.M112.424077

De Vienne, D. M. (2016). Lifemap: Exploring the Entire Tree of Life. PLOS Biology, 14(12), e2001624. 10.1371/journal.pbio.2001624

Eswaramoorthy, S., Bonanno, J. B., Burley, S. K., & Swaminathan, S. (2006). Mechanism of action of a flavin-containing monooxygenase. Proceedings of the National Academy of Sciences, 103(26), 9832–9837. 10.1073/pnas.0602398103

Florean, M., Luck, K., Hong, B., Nakamura, Y., O’Connor, S. E., & Köllner, T. G. (2023). Reinventing metabolic pathways: Independent evolution of benzoxazinoids in flowering plants. Proceedings of the National Academy of Sciences, 120(42), e2307981120. 10.1073/pnas.2307981120

Florean, M., Schultz, H., Wurlitzer, J., O’Connor, S. E., & Köllner, T. G. (2025). Independent evolution of plant natural products: Formation of benzoxazinoids in Consolida orientalis (Ranunculaceae). Journal of Biological Chemistry, 301(1), 108019. 10.1016/j.jbc.2024.108019

Fraaije, M. W., Kamerbeek, N. M., Van Berkel, W. J. H., & Janssen, D. B. (2002). Identification of a Baeyer–Villiger monooxygenase sequence motif. FEBS Letters, 518(1–3), 43–47. 10.1016/S0014-5793(02)02623-6

Goodstein, D. M., Shu, S., Howson, R., Neupane, R., Hayes, R. D., Fazo, J., Mitros, T., Dirks, W., Hellsten, U., Putnam, N., & Rokhsar, D. S. (2012). Phytozome: A comparative platform for green plant genomics. Nucleic Acids Research, 40(D1), D1178–D1186. 10.1093/nar/gkr944

Greene, L. H., Lewis, T. E., Addou, S., Cuff, A., Dallman, T., Dibley, M., Redfern, O., Pearl, F., Nambudiry, R., Reid, A., Sillitoe, I., Yeats, C., Thornton, J. M., & Orengo, C. A. (2007). The CATH domain structure database: New protocols and classification levels give a more comprehensive resource for exploring evolution. Nucleic Acids Research, 35(Database), D291–D297. 10.1093/nar/gkl959

Grigoriev, I. V., Hayes, R. D., Calhoun, S., Kamel, B., Wang, A., Ahrendt, S., Dusheyko, S., Nikitin, R., Mondo, S. J., Salamov, A., Shabalov, I., & Kuo, A. (2021). PhycoCosm, a comparative algal genomics resource. Nucleic Acids Research, 49(D1), D1004–D1011. 10.1093/nar/gkaa898

Hansen, B. G., Kliebenstein, D. J., & Halkier, B. A. (2007). Identification of a flavin-monooxygenase as the S -oxygenating enzyme in aliphatic glucosinolate biosynthesis in Arabidopsis. The Plant Journal, 50(5), 902–910. 10.1111/j.1365-313X.2007.03101.x

Hartmann, M., Zeier, T., Bernsdorff, F., Reichel-Deland, V., Kim, D., Hohmann, M., Scholten, N., Schuck, S., Bräutigam, A., Hölzel, T., Ganter, C., & Zeier, J. (2018). Flavin Monooxygenase-Generated N-Hydroxypipecolic Acid Is a Critical Element of Plant Systemic Immunity. Cell, 173(2), 456-469.e16. 10.1016/j.cell.2018.02.049

Healey, A. L., Piatkowski, B., Lovell, J. T., Sreedasyam, A., Carey, S. B., Mamidi, S., Shu, S., Plott, C., Jenkins, J., Lawrence, T., Aguero, B., Carrell, A. A., Nieto-Lugilde, M., Talag, J., Duffy, A., Jawdy, S., Carter, K. R., Boston, L.-B., Jones, T., … Shaw, A. J. (2023). Newly identified sex chromosomes in the Sphagnum (peat moss) genome alter carbon sequestration and ecosystem dynamics. Nature Plants, 9(2), 238–254. 10.1038/s41477-022-01333-5

Holmes, E. C., Chen, Y.-C., Sattely, E. S., & Mudgett, M. B. (2019). An engineered pathway for N - hydroxy-pipecolic acid synthesis enhances systemic acquired resistance in tomato. Science Signaling, 12(604), eaay3066. 10.1126/scisignal.aay3066

Huijbers, M. M., Montersino, S., Westphal, A. H., Tischler, D., & van Berkel, W. J. (2014). Flavin dependent monooxygenases. Arch Biochem Biophys, 544, 2–17. 10.1016/j.abb.2013.12.005

Jiang, K., Møller, B. L., Luo, S., Yang, Y., Nelson, D. R., Jakobsen Neilson, E. H., Christensen, J. M., Hua, K., Hu, C., Zeng, X., Motawie, M. S., Wan, T., Hu, G.-W., Onjalalaina, G. E., Wang, Y., Gaitán-Espitia, J. D., Wang, Z., Xu, X.-Y., He, J., … Huang, W.-C. (2025). Genomic, transcriptomic, and metabolomic analyses reveal convergent evolution of oxime biosynthesis in Darwin’s orchid. Molecular Plant, 18(3), 392–415. 10.1016/j.molp.2024.12.010

Katoh, K., & Standley, D. M. (2013). MAFFT Multiple Sequence Alignment Software Version 7: Improvements in Performance and Usability. Molecular Biology and Evolution, 30(4), 772–780. 10.1093/molbev/mst010

Kong, W., Li, J., Yu, Q., Cang, W., Xu, R., Wang, Y., & Ji, W. (2016). Two Novel Flavin-Containing Monooxygenases Involved in Biosynthesis of Aliphatic Glucosinolates. Frontiers in Plant Science, 7. 10.3389/fpls.2016.01292

Li, F.-W., Brouwer, P., Carretero-Paulet, L., Cheng, S., de Vries, J., Delaux, P.-M., Eily, A., Koppers, N., Kuo, L.-Y., Li, Z., Simenc, M., Small, I., Wafula, E., Angarita, S., Barker, M. S., Bräutigam, A., dePamphilis, C., Gould, S., Hosmani, P. S., … Pryer, K. M. (2018). Fern genomes elucidate land plant evolution and cyanobacterial symbioses. Nature Plants, 4(7), 460–472. 10.1038/s41477-018-0188-8

Liscombe, D. K., Kamiyoshihara, Y., Ghironzi, J., Kempthorne, C. J., Hooton, K., Bulot, B., Kanellis, V., McNulty, J., Lam, N. B., Nadeau, L. F., Pautler, M., Tieman, D. M., Klee, H. J., & Goulet, C. (2022). A flavin-dependent monooxygenase produces nitrogenous tomato aroma volatiles using cysteine as a nitrogen source. Proceedings of the National Academy of Sciences, 119(7), e2118676119. 10.1073/pnas.2118676119

Mascotti, M. L., Juri Ayub, M., Furnham, N., Thornton, J. M., & Laskowski, R. A. (2016). Chopping and Changing: The Evolution of the Flavin-dependent Monooxygenases. Journal of Molecular Biology, 428(15), 3131–3146. 10.1016/j.jmb.2016.07.003

Mashiguchi, K., Tanaka, K., Sakai, T., Sugawara, S., Kawaide, H., Natsume, M., Hanada, A., Yaeno, T., Shirasu, K., Yao, H., McSteen, P., Zhao, Y., Hayashi, K., Kamiya, Y., & Kasahara, H. (2011). The main auxin biosynthesis pathway in Arabidopsis. Proceedings of the National Academy of Sciences, 108(45), 18512–18517. 10.1073/pnas.1108434108

Meng, E. C., Goddard, T. D., Pettersen, E. F., Couch, G. S., Pearson, Z. J., Morris, J. H., & Ferrin, T. E. (2023). UCSF CHIMERAX: Tools for structure building and analysis. Protein Science, 32(11), e4792. 10.1002/pro.4792

Meng, E. C., Pettersen, E. F., Couch, G. S., Huang, C. C., & Ferrin, T. E. (2006). Tools for integrated sequence-structure analysis with UCSF Chimera. BMC Bioinformatics, 7(1), 339. 10.1186/1471-2105-7-339

Nicoll, C. R., & Mascotti, M. L. (2023). Investigating the biochemical signatures and physiological roles of the FMO family using molecular phylogeny. BBA Advances, 4, 100108. 10.1016/j.bbadva.2023.100108

One Thousand Plant Transcriptomes Initiative. (2019). One thousand plant transcriptomes and the phylogenomics of green plants. Nature, 574(7780), 679–685. 10.1038/s41586-019-1693-2

Orengo, C., Michie, A., Jones, S., Jones, D., Swindells, M., & Thornton, J. (1997). CATH – a hierarchic classification of protein domain structures. Structure, 5(8), 1093–1109. 10.1016/S0969-2126(97)00260-8

Perillo, B., Tramontano, A., Pezone, A., & Migliaccio, A. (2020). LSD1: More than demethylation of histone lysine residues. Experimental & Molecular Medicine, 52(12), 1936–1947. 10.1038/s12276-020-00542-2

Sánchez-Pérez, R., & Neilson, E. Hj. (2024). The case for sporadic cyanogenic glycoside evolution in plants. Current Opinion in Plant Biology, 81, 102608. 10.1016/j.pbi.2024.102608

Stehr, M., Diekmann, H., Smau, L., Seth, O., Ghisla, S., Singh, M., & Macheroux, P. (1998). A hydrophobic sequence motif common to N-hydroxylating enzymes. Trends in Biochemical Sciences, 23(2), 56–57. 10.1016/S0968-0004(97)01166-3

Sundell, D., Mannapperuma, C., Netotea, S., Delhomme, N., Lin, Y., Sjödin, A., Van De Peer, Y., Jansson, S., Hvidsten, T. R., & Street, N. R. (2015). The Plant Genome Integrative Explorer Resource: PlantGen IE .org. New Phytologist, 208(4), 1149–1156. 10.1111/nph.13557

Tan, Q. W., Lim, P. K., Chen, Z., Pasha, A., Provart, N., Arend, M., Nikoloski, Z., & Mutwil, M. (2023). Cross-stress gene expression atlas of Marchantia polymorpha reveals the hierarchy and regulatory principles of abiotic stress responses. Nature Communications, 14(1), 986. 10.1038/s41467-023-36517-w

Thodberg, S., & Jakobsen Neilson, E. H. (2020). The “Green” FMOs: Diversity, Functionality and Application of Plant Flavoproteins. Catalysts, 10(3), 329. 10.3390/catal10030329

Thodberg, S., Sørensen, M., Bellucci, M., Crocoll, C., Bendtsen, A. K., Nelson, D. R., Motawia, M. S., Møller, B. L., & Neilson, E. H. J. (2020). A flavin-dependent monooxygenase catalyzes the initial step in cyanogenic glycoside synthesis in ferns. Communications Biology, 3(1), 507. 10.1038/s42003-020-01224-5

Valentino, H., Campbell, A. C., Schuermann, J. P., Sultana, N., Nam, H. G., LeBlanc, S., Tanner, J. J., & Sobrado, P. (2020). Structure and function of a flavin-dependent S-monooxygenase from garlic (Allium sativum). Journal of Biological Chemistry, 295(32), 11042–11055. 10.1074/jbc.RA120.014484

Van Den Heuvel, R. H. H., Curti, B., Vanoni, M. A., & Mattevi, A. (2004). Glutamate synthase: A fascinating pathway from L-glutamine to L-glutamate. Cellular and Molecular Life Sciences (CMLS), 61(6), 669–681. 10.1007/s00018-003-3316-0

Van Kempen, M., Kim, S. S., Tumescheit, C., Mirdita, M., Lee, J., Gilchrist, C. L. M., Söding, J., & Steinegger, M. (2024). Fast and accurate protein structure search with Foldseek. Nature Biotechnology, 42(2), 243–246. 10.1038/s41587-023-01773-0

Wang, H., Yang, Q., Tan, S., Wang, T., Zhang, Y., Yang, Y., Yin, W., Xia, X., Guo, H., & Li, Z. (2022). Regulation of cytokinin biosynthesis using PtRD26pro -IPT module improves drought tolerance through PtARR10-PtYUC4/5-mediated reactive oxygen species removal in Populus. Journal of Integrative Plant Biology, 64(3), 771–786. 10.1111/jipb.13218

Wang, L., Zhou, Y., Ding, Y., Chen, C., Chen, X., Su, N., Zhang, X., Pan, Y., & Li, J. (2023). Novel flavin-containing monooxygenase protein FMO1 interacts with CAT2 to negatively regulate drought tolerance through ROS homeostasis and ABA signaling pathway in tomato. Horticulture Research, 10(4), uhad037. 10.1093/hr/uhad037

Won, C., Shen, X., Mashiguchi, K., Zheng, Z., Dai, X., Cheng, Y., Kasahara, H., Kamiya, Y., Chory, J., & Zhao, Y. (2011). Conversion of tryptophan to indole-3-acetic acid by TRYPTOPHAN AMINOTRANSFERASES OF ARABIDOPSIS and YUCCAs in Arabidopsis. Proceedings of the National Academy of Sciences, 108(45), 18518–18523. 10.1073/pnas.1108436108

Zhao, Y., Christensen, S. K., Fankhauser, C., Cashman, J. R., Cohen, J. D., Weigel, D., & Chory, J. (2001). A Role for Flavin Monooxygenase-Like Enzymes in Auxin Biosynthesis. Science, 291(5502), 306–309. 10.1126/science.291.5502.306

Ziegler, D. M. (1988). Flavin-containing monooxygenases: Catalytic mechanism and substrate specificities. Drug Metabolism Reviews, 19(1), 1–32. 10.3109/03602538809049617

