## Supplementary Information for "A plant-centric investigation of Class B Flavin-dependent Monooxygenase evolution and structural diversity"

Supplementary Figure 1: Subtree of streptophyte species included this study overlaid the entire taxonomy of streptophyta. Snapshot from Lifemap server (De Vienne, 2016)

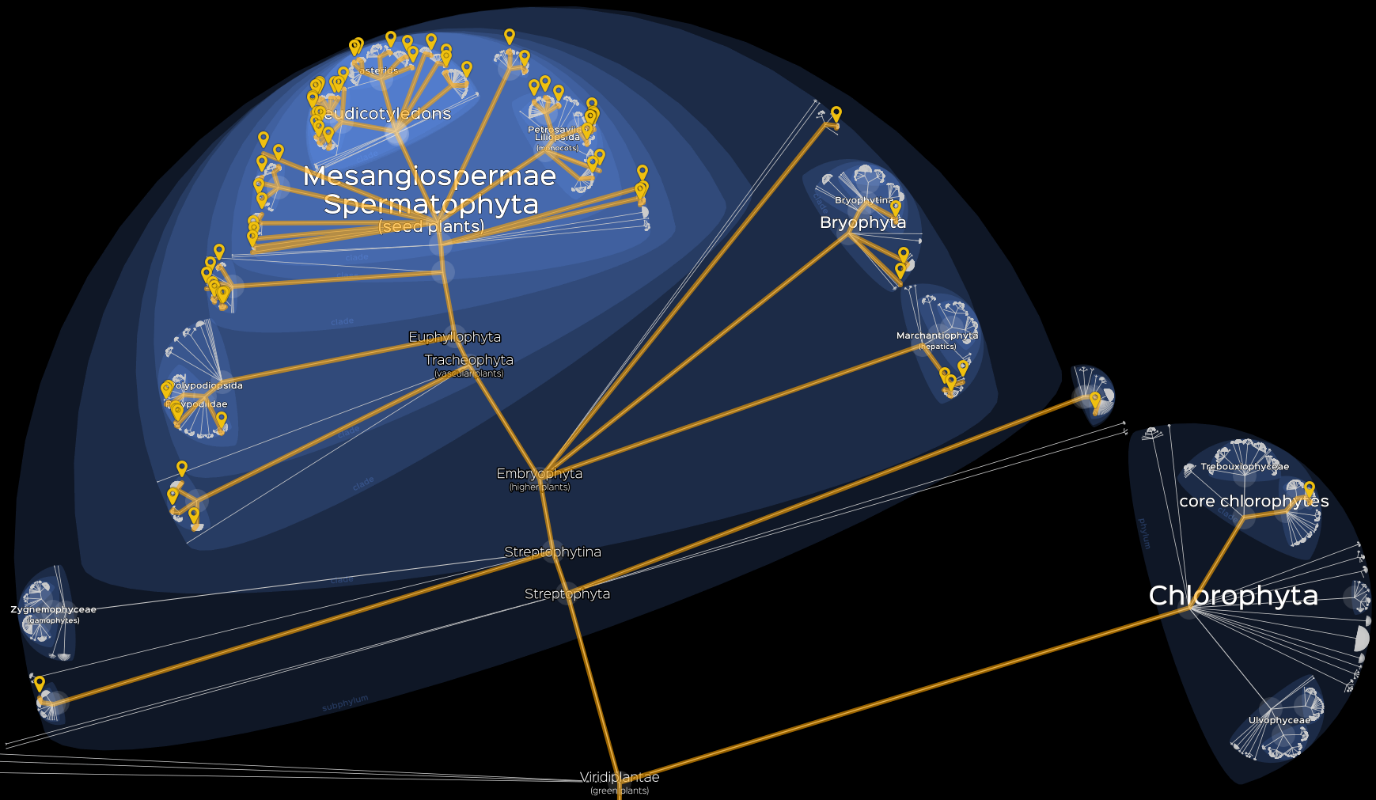

**Supplementary Figure 2: Alphafold 3 predicted structure of AtFMO1 superimposed on AtYUC10 to assign Class B FMO motif; FMO-ID (purple), NADPH-binding (orange) and ATG (aquamarine) motifs of AtFMO1 (colored in white).**

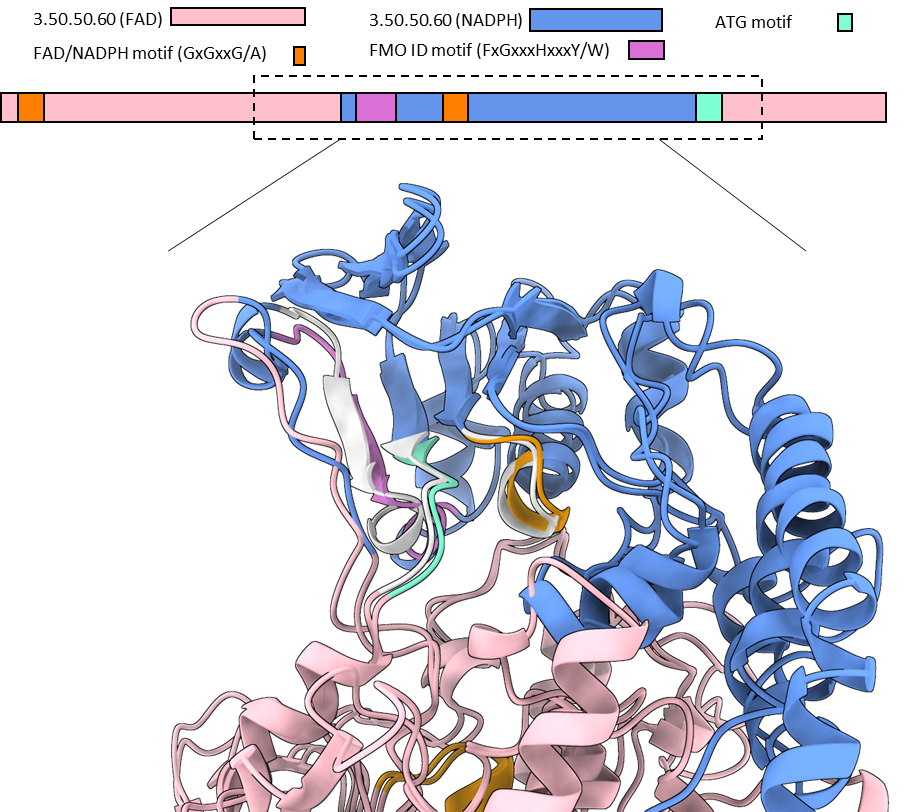

**Supplementary Figure 3: Gene expression of two Marchantia SeedLess FMOs visualized in the Marchantia Atlas eFP Browser (Tan et al., 2023).**

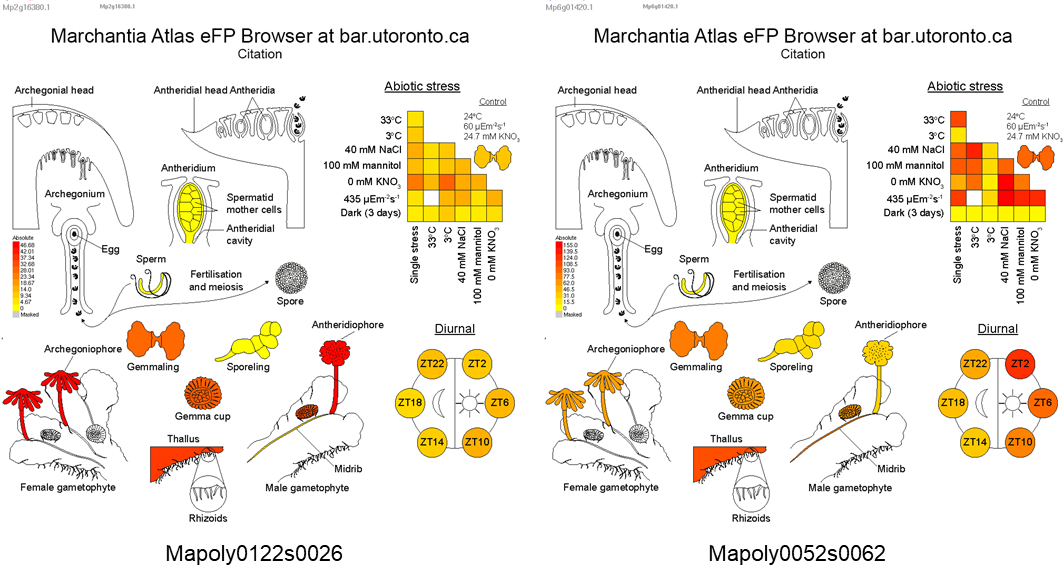

**Supplementary Table 1 BLASTP queries for sequence acquisition**

| **Species** | **GenomeID** | **Lineage** | **Accession** | **FMO Family** |
| --- | --- | --- | --- | --- |
| Marchantia polymorpha | Mpolymorpha_v3.1 | Liverwort | Mapoly0063s0040 | YUCCA |
| Physcomitrium patens | Ppatens_v6.1 | Moss | Pp3c1_11500V3 | YUCCA |
| Selaginella moellendorffii | Smoellendorffii_v1.0 | Lycophyte | 75206 | YUCCA |
| Ceratopteris richardii | Crichardii_v2.1 | Monilophyte (fern) | Ceric.05G083300 | YUCCA |
| Hordeum vulgare | Hvulgare_r1 | Monocot | HORVU7Hr1G013170 | YUCCA |
| Arabidopsis thaliana | Athaliana_Araport11 | Eudicot (Rosid) | AT5G25620 | YUCCA |
| Physcomitrium patens | Ppatens_v6.1 | Moss | Pp6c18_10300V6 | BVMO |
| Chlamydomonas reinhardtii | Creinhardtii_v5.6 | Chlorophyte | Cre03.g167150.t1.2 | S-Ox |
| Marchantia polymorpha | Mpolymorpha_v3.1 | Liverwort | Mapoly0002s0275 | S-Ox |
| Physcomitrium patens | Ppatens_v6.1 | Moss | Pp3c11_14620V3 | S-Ox |
| Selaginella moellendorffii | Smoellendorffii_v1.0 | Lycophyte | 413958 | S-Ox |
| Ceratopteris richardii | Crichardii_v2.1 | Monilophyte (fern) | Ceric.34G032600 | S-Ox |
| Hordeum vulgare | Hvulgare_r1 | Monocot | HORVU1Hr1G053900 | S-Ox |
| Arabidopsis thaliana | Athaliana_Araport11 | Eudicot (Rosid) | AT1G65860 | S-Ox |
| Selaginella moellendorffii | Smoellendorffii_v1.0 | Lycophyte | 266739 | N-Ox |
| Ceratopteris richardii | Crichardii_v2.1 | Monilophyte (fern) | Ceric.23G070500 | N-Ox |
| Picea abies | Picea_abies | Gymnosperm | MA_10430096g0010 | N-Ox |
| Hordeum vulgare | Hvulgare_r1 | Monocot | HORVU7Hr1G121260 | N-Ox |
| Arabidopsis thaliana | Athaliana_Araport11 | Eudicot (Rosid) | AT1G19250 | N-Ox |

**Supplementary Table 2 Species List with data source**

| **Species** | **Genome ID:** | **NCBI (txid)** | **Common Name** | **Lineage** | **Source** | **BLAST source** |
| --- | --- | --- | --- | --- | --- | --- |
| Acorus americanus | Aamericanus_v1.1 | 263995 | American sweet flag | Monocot | Phytozome | Online |
| Aquilegia coerulea | Acoerulea_v3.1 | 218851 | blue columbine | Basal Eudicots+ | Phytozome | Online |
| Ananas comosus | Acomosus_v3 | 4615 | Pineapple | Monocot | Phytozome | Online |
| Adiantum capillus-veneris | Adiantum_capillus-veneris | 13818 | Southern maidenhair fern | Monilophyte | FernBase | Local |
| Allium sativum | Allium_sativum | 4682 | Garlic | Monocot | (Sun et al., 2020) | Local |
| Alsophila Spinulosa | Alsophila_spinulosa | 204586 | Flying spider-monkey tree fern | Monilophyte | (Huang et al., 2022) | Local |
| Anthoceros angustus | Anthoceros_angustus | 48387 | (hornwort) | Hornwort | (Zhang et al., 2020) | Local |
| Asparagus officinalis | Aofficinalis_V1.1 | 4686 | Asparagus | Monocot | Phytozome | Online |
| Arabidopsis thaliana | Athaliana_Araport11 | 3702 | Thale cress | Rosids | Phytozome | Online |
| Amborella trichopoda | Atrichopoda_v1.0 | 13333 | Amboralla | ANA Grade Angiosperms | Phytozome | Online |
| Azolla filiculoides | Azolla_filiculoides | 84609 | Mosquito fern | Monilophyte | FernBase | Local |
| Brachypodium distachyon | Bdistachyon_v3.2 | 15368 | Purple false brome | Monocot | Phytozome | Online |
| Brassica rapa | BrapaFPsc_v1.3 | 3711 | Rape / field mustard | Rosids | Phytozome | Online |
| Buxus sinica | Buxus_sinica | 153572 | Korean littleleaf boxwood | Basal Eudicots+ | (Chanderbali et al., 2022) | Local |
| Coffea arabica | Carabica_v0.5 | 13443 | Coffee | Asterids | Phytozome | Online |
| Ceratophyllum demersum | Ceratophyllum_demersum | 4428 | Rigid hornwort | Basal Eudicots+ | (Yang et al., 2020) | Local |
| Ceratopteris richardii | Ceratopteris_richardii | 49495 | Triangle waterfern | Monilophyte | FernBase | Local |
| Chara Braunii | Chara_Braunii | 69332 | (Green Algea) | Charophyta | phycocosm | Online |
| Chloranthus spicatus | Chloranthus_spicatus | 13006 | Charan | Basal Eudicots+ | (Guo et al., 2021) | Local |
| Cinnamomum kanehirae | Ckanehirae_v3 | 337451 | Stout camphor tree | Magnoliids+ | Phytozome | Online |
| Carica papaya | Cpapaya_ASGPBv0.4 | 3649 | Papaya / PawPaw | Rosids | Phytozome | Online |
| Chenopodium quinoa | Cquinoa_v1.0 | 63459 | Quinoa | Basal Eudicots+ | Phytozome | Online |
| Chlamydomonas reinhardtii | Creinhardtii_CC-4532_v6.1 | 3055 | (Green Algea) | Chlorophyte | Phytozome | Online |
| Cucumis sativus | Csativus_v1.0 | 3659 | Cucumber | Rosids | Phytozome | Online |
| Citrus sinensis | Csinensis_v1.1 | 2711 | Orange | Rosids | Phytozome | Online |
| Cycas panzhihuaensis | Cycas_panzhihuaensis | 123604 | Dukou sago palm | Gymnosperm | (Liu et al., 2022) | Local |
| Daucus carota | Dcarota_subsp_sativus_v3.0 | 4039 | Carrot | Asterids | Phytozome | Online |
| Diphasiastrum complanatum | Dcomplanatum_v3.1 | 34168 | Groundcedar | Lycophyte | Phytozome | Online |
| Eleusine coracana | Ecoracana_v1.1 | 4511 | Finger millet | Monocot | Phytozome | Online |
| Eucalyptus grandis | Egrandis_v2.0 | 71139 | Flooded gum | Rosids | Phytozome | Online |
| Euryale ferox | Euryale_ferox | 4414 | Prickly waterlily | ANA Grade Angiosperms | (Yang et al., 2020) | Local |
| Fragaria vesca | Fvesca_v4.0.a2 | 57918 | Wild strawberry | Rosids | Phytozome | Online |
| Gossypium hirsutum | Ghirsutum_v3.1 | 3635 | Upland cotton | Rosids | Phytozome | Online |
| Ginkgo biloba | Ginkgo_biloba | 3311 | Ginkgo | Gymnosperm | PlantGenie | Online |
| Glycine Max | Gmax_Wm82.a6.v1 | 3847 | Soybean | Rosids | Phytozome | Online |
| Gnetum montanum | Gnetum_montanum | 3381 | Gam nui | Gymnosperm | PlantGenie | Online |
| Hydrangea quercifolia | Hquercifolia_v1.1 | 60124 | Oakleaf hydrangea | Asterids | Phytozome | Online |
| Hordeum vulgare | Hvulgare_Morex_V3 | 4513 | Barley | Monocot | Phytozome | Online |
| Isoetes sinensis | Isoetes_sinensis | 283158 | Narrow quillwort | Lycophyte | GigaDB | Local |
| Kalanchoe laxiflora | Klaxiflora_v1.1 | 1670617 | Milky Widow's Tears | Basal Eudicots+ | Phytozome | Online |
| Klebsormidium nitens | Klebsormidium_nitens | 105231 | (Green Algea) | Charophyta | Phycocosm | Online |
| Liriodendron tulipifera | Ltulipifera_YP108A_v1.1 | 3415 | Yellow poplar / tulip tree | Magnoliids+ | Phytozome | Online |
| Lunularia cruciata | Lunularia_cruciata | 56931 | Crescent-cup liverwort | Liverwort | (Linde et al., 2021) | Local |
| Lycopodium clavatum | Lycopodium_clavatum | 3252 | Common club moss | Lycophyte | (Yu et al., 2023) | Local |
| Macadamia integrifolia | Macadamia_integrifolia | 60698 | Macadamia Nut | Basal Eudicots+ | (Lin et al., 2022) | Local |
| Musa acuminata | Macuminata_v1 | 4641 | Dwarf Cavendish Banana | Monocot | Phytozome | Online |
| Marsilea vestita | Marsilea_vestita | 59764 | Hairy Water Clover (Fern) | Monilophyte | FernBase | Local |
| Marchantia polymorpha | Mpolymorpha_v3.1 | 3197 | Common liverwort | Liverwort | Phytozome | Online |
| Nymphaea colorata | Ncolorata_v1.2 | 210225 | Blue-Petal Water Lily | ANA Grade Angiosperms | Phytozome | Online |
| Oryza sativa | Osativa_v7.0 | 39947 | Rice | Monocot | Phytozome | Online |
| Papaver somniferum | Papaver_somniferum | 3469 | Opium poppy | Basal Eudicots+ | Papaver genomics | Local |
| Phalaenopsis equestris | Phalaenopsis_equestris | 78828 | Darwins Orchid | Monocot | PRJNA192198 | Local /NCBI |
| Picea abies | Picea_abies | 3329 | Spruce | Gymnosperm | PlantGenie | Online |
| Pinus taeda | Pinus_taeda | 3352 | Loblolly pine | Gymnosperm | PlantGenie | Online |
| Piper nigrum | Piper_nigrum | 13216 | Black Pepper | Magnoliids+ | (L. Hu et al., 2019) | Local |
| Pisum Sativum | Pisum_sativum | 3888 | Common Pea | Rosids | (Kreplak et al., 2019) | Local |
| Physcomitrium patens | Ppatens_v6.1 | 3218 | Spreading earthmoss | Moss | Phytozome | Online |
| Prunus persica | Ppersica_v2.1 | 3760 | Peach | Rosids | Phytozome | Online |
| Pseudotsuga menziesii | Pseudotsuga_menziesii | 3357 | Douglas fir | Gymnosperm | PlantGenie | Online |
| Populus trichocarpa | Ptrichocarpa_v4.1 | 3694 | Black cottonwood | Rosids | Phytozome | Online |
| Quercus robur | Quercus_robur | 38942 | Common Oak | Rosids | (Plomion et al., 2018) | Local |
| Rhododendron delavayi | Rhododendron_delavayi | 321363 | Magnolia | Asterids | (Wu et al., 2023) | Local |
| Ricciocarpos natans | Ricciocarpos_natans | 53035 | Fringed heartwort | Liverwort | (Linde et al., 2021) | Local |
| Salvinia cucullata | Salvinia_cucullata | 32188 | Asian Watermoss | Monilophyte | FernBase | Local |
| Santalum yasi | Santalum_yasi | 453089 | Fijian Sandalwood | Basal Eudicots+ | (Hong et al., 2023) | Local |
| Sorghum bicolor | Sbicolor_v5.1 | 4558 | Sorghum / Durra | Monocot | Phytozome | Online |
| Sphagnum fallax | Sfallax_v1.1 | 53036 | Flat-topped bogmoss | Moss | Phytozome | Online |
| Solanum lycopersicum | Slycopersicum_ITAG5.0 | 4081 | Tomato | Asterids | Phytozome | Online |
| Selaginella moellendorffii | Smoellendorffii_v1.0 | 88036 | Gemmiferous Spikemoss | Lycophyte | Phytozome | Online |
| Saponaria officinalis | Sofficinalis_v1.1 | 3572 | Common soapwort | Basal Eudicots+ | Phytozome | Online |
| Solanum tuberosum | Stuberosum_v6.1 | 4113 | Potato | Asterids | Phytozome | Online |
| Takakia lepidozioides | Takakia_lepidozioides | 37425 | Takakia | Moss | (R. Hu et al., 2023) | Local |
| Taxus chinensis | Taxus_chinensis | 29808 | Chinese Yew | Gymnosperm | (Xiong et al., 2021) | Local |
| Tetracentron sinense | Tetracentron_sinense | 13715 | Spur-leaf | Basal Eudicots+ | (Chanderbali et al., 2022) | Local |
| Thuja plicata | Tplicata_v3.1 | 3316 | Western red cedar | Gymnosperm | Phytozome | Online |
| Vitis vinifera | Vvinifera_v2.1 | 29760 | European wine grape | Rosids | Phytozome | Online |
| Zostera Marina | Zmarina_v3.1 | 29655 | Common eelgrass | Monocot | Phytozome | Online |
| Zea mays | Zmays_RefGen_V4 | 4577 | Maize (Corn) | Monocot | Phytozome | Online |

Supplementary Table 3: RMSD values for comparison between modeled protein structures, accessions can be found in methods

| AsFMO1 | AsFMO1 | |  |  |  |  |  |  |  |
| --- | --- | --- | --- | --- | --- | --- | --- | --- | --- |
| AtFMO1 | 10.29 | AtFMO1 | |  |  |  |  |  |  |
| AtGS-OX1 | 0.905 | 10.027 | AtGS-OX1 | |  |  |  |  |  |
| AtYUC10 | 7.243 | 6.87 | 6.647 | AtYUC10 | |  |  |  |  |
| SfS-OX-like 2 | 8.734 | 10.871 | 9.676 | 6.738 | SfS-OX-like 2 | |  |  |  |
| MpS-Ox-like 1 | 4.334 | 6.891 | 5.793 | 5.273 | 9.979 | MpS-Ox-like 1 | |  |  |
| PpBVMO | 17.191 | 10.952 | 18.733 | 8.266 | 8.236 | 15.23 | PpBVMO | |  |
| ScBVMO | 9.876 | 9.61 | 10.99 | 8.52 | 10.391 | 15.019 | 5.112 | ScBVMO | |
| SfN-Ox-like | 7.833 | 6.105 | 8.514 | 5.564 | 10.013 | 8.708 | 10.206 | 9.576 | SfN-Ox-like |
| CrSeedless FMO | 10.811 | 6.823 | 11.79 | 5.268 | 21.365 | 6.329 | 9.308 | 9.098 | 9.403 |

**References**

Chanderbali, A. S., Jin, L., Xu, Q., Zhang, Y., Zhang, J., Jian, S., Carroll, E., Sankoff, D., Albert, V. A., Howarth, D. G., Soltis, D. E., & Soltis, P. S. (2022). Buxus and Tetracentron genomes help resolve eudicot genome history. *Nature Communications*, *13*(1), 643. https://doi.org/10.1038/s41467-022-28312-w

Guo, X., Fang, D., Sahu, S. K., Yang, S., Guang, X., Folk, R., Smith, S. A., Chanderbali, A. S., Chen, S., Liu, M., Yang, T., Zhang, S., Liu, X., Xu, X., Soltis, P. S., Soltis, D. E., & Liu, H. (2021). Chloranthus genome provides insights into the early diversification of angiosperms. *Nature Communications*, *12*(1), 6930. https://doi.org/10.1038/s41467-021-26922-4

Hong, Z., Peng, D., Tembrock, L. R., Liao, X., Xu, D., Liu, X., & Wu, Z. (2023). Chromosome-level genome assemblies from two sandalwood species provide insights into the evolution of the Santalales. *Communications Biology*, *6*(1), 587. https://doi.org/10.1038/s42003-023-04980-2

Hu, L., Xu, Z., Wang, M., Fan, R., Yuan, D., Wu, B., Wu, H., Qin, X., Yan, L., Tan, L., Sim, S., Li, W., Saski, C. A., Daniell, H., Wendel, J. F., Lindsey, K., Zhang, X., Hao, C., & Jin, S. (2019). The chromosome-scale reference genome of black pepper provides insight into piperine biosynthesis. *Nature Communications*, *10*(1), 4702. https://doi.org/10.1038/s41467-019-12607-6

Hu, R., Li, X., Hu, Y., Zhang, R., Lv, Q., Zhang, M., Sheng, X., Zhao, F., Chen, Z., Ding, Y., Yuan, H., Wu, X., Xing, S., Yan, X., Bao, F., Wan, P., Xiao, L., Wang, X., Xiao, W., … He, Y. (2023). Adaptive evolution of the enigmatic Takakia now facing climate change in Tibet. *Cell*, *186*(17), 3558-3576.e17. https://doi.org/10.1016/j.cell.2023.07.003

Huang, X., Wang, W., Gong, T., Wickell, D., Kuo, L.-Y., Zhang, X., Wen, J., Kim, H., Lu, F., Zhao, H., Chen, S., Li, H., Wu, W., Yu, C., Chen, S., Fan, W., Chen, S., Bao, X., Li, L., … Li, Q. (2022). The flying spider-monkey tree fern genome provides insights into fern evolution and arborescence. *Nature Plants*, *8*(5), 500–512. https://doi.org/10.1038/s41477-022-01146-6

Kreplak, J., Madoui, M.-A., Cápal, P., Novák, P., Labadie, K., Aubert, G., Bayer, P. E., Gali, K. K., Syme, R. A., Main, D., Klein, A., Bérard, A., Vrbová, I., Fournier, C., d’Agata, L., Belser, C., Berrabah, W., Toegelová, H., Milec, Z., … Burstin, J. (2019). A reference genome for pea provides insight into legume genome evolution. *Nature Genetics*, *51*(9), 1411–1422. https://doi.org/10.1038/s41588-019-0480-1

Lin, J., Zhang, W., Zhang, X., Ma, X., Zhang, S., Chen, S., Wang, Y., Jia, H., Liao, Z., Lin, J., Zhu, M., Xu, X., Cai, M., Zeng, H., Wan, J., Yang, W., Matsumoto, T., Hardner, C., Nock, C. J., & Ming, R. (2022). Signatures of selection in recently domesticated macadamia. *Nature Communications*, *13*(1), 242. https://doi.org/10.1038/s41467-021-27937-7

Linde, A.-M., Eklund, D. M., Cronberg, N., Bowman, J. L., & Lagercrantz, U. (2021). Rates and patterns of molecular evolution in bryophyte genomes, with focus on complex thalloid liverworts, Marchantiopsida. *Molecular Phylogenetics and Evolution*, *165*, 107295. https://doi.org/10.1016/j.ympev.2021.107295

Liu, Y., Wang, S., Li, L., Yang, T., Dong, S., Wei, T., Wu, S., Liu, Y., Gong, Y., Feng, X., Ma, J., Chang, G., Huang, J., Yang, Y., Wang, H., Liu, M., Xu, Y., Liang, H., Yu, J., … Zhang, S. (2022). The Cycas genome and the early evolution of seed plants. *Nature Plants*, *8*(4), 389–401. https://doi.org/10.1038/s41477-022-01129-7

Plomion, C., Aury, J.-M., Amselem, J., Leroy, T., Murat, F., Duplessis, S., Faye, S., Francillonne, N., Labadie, K., Le Provost, G., Lesur, I., Bartholomé, J., Faivre-Rampant, P., Kohler, A., Leplé, J.-C., Chantret, N., Chen, J., Diévart, A., Alaeitabar, T., … Salse, J. (2018). Oak genome reveals facets of long lifespan. *Nature Plants*, *4*(7), 440–452. https://doi.org/10.1038/s41477-018-0172-3

Sun, X., Zhu, S., Li, N., Cheng, Y., Zhao, J., Qiao, X., Lu, L., Liu, S., Wang, Y., Liu, C., Li, B., Guo, W., Gao, S., Yang, Z., Li, F., Zeng, Z., Tang, Q., Pan, Y., Guan, M., … Liu, T. (2020). A Chromosome-Level Genome Assembly of Garlic (Allium sativum) Provides Insights into Genome Evolution and Allicin Biosynthesis. *Molecular Plant*, *13*(9), 1328–1339. https://doi.org/10.1016/j.molp.2020.07.019

Wu, X., Zhang, L., Wang, X., Zhang, R., Jin, G., Hu, Y., Yang, H., Wu, Z., Ma, Y., Zhang, C., & Wang, J. (2023). Evolutionary history of two evergreen Rhododendron species as revealed by chromosome-level genome assembly. *Frontiers in Plant Science*, *14*. https://doi.org/10.3389/fpls.2023.1123707

Xiong, X., Gou, J., Liao, Q., Li, Y., Zhou, Q., Bi, G., Li, C., Du, R., Wang, X., Sun, T., Guo, L., Liang, H., Lu, P., Wu, Y., Zhang, Z., Ro, D.-K., Shang, Y., Huang, S., & Yan, J. (2021). The Taxus genome provides insights into paclitaxel biosynthesis. *Nature Plants*, *7*(8), 1026–1036. https://doi.org/10.1038/s41477-021-00963-5

Yang, Y., Sun, P., Lv, L., Wang, D., Ru, D., Li, Y., Ma, T., Zhang, L., Shen, X., Meng, F., Jiao, B., Shan, L., Liu, M., Wang, Q., Qin, Z., Xi, Z., Wang, X., Davis, C. C., & Liu, J. (2020). Prickly waterlily and rigid hornwort genomes shed light on early angiosperm evolution. *Nature Plants*, *6*(3), 215–222. https://doi.org/10.1038/s41477-020-0594-6

Yu, J.-G., Tang, J.-Y., Wei, R., Lan, M.-F., Xiang, R.-C., Zhang, X.-C., & Xiang, Q.-P. (2023). The first homosporous lycophyte genome revealed the association between the recent dynamic accumulation of LTR-RTs and genome size variation. *Plant Molecular Biology*, *112*(6), 325–340. https://doi.org/10.1007/s11103-023-01366-0

Zhang, J., Fu, X.-X., Li, R.-Q., Zhao, X., Liu, Y., Li, M.-H., Zwaenepoel, A., Ma, H., Goffinet, B., Guan, Y.-L., Xue, J.-Y., Liao, Y.-Y., Wang, Q.-F., Wang, Q.-H., Wang, J.-Y., Zhang, G.-Q., Wang, Z.-W., Jia, Y., Wang, M.-Z., … Chen, Z.-D. (2020). The hornwort genome and early land plant evolution. *Nature Plants*, *6*(2), 107–118. https://doi.org/10.1038/s41477-019-0588-4
